# Nestin selectively facilitates the phosphorylation of the Lissencephaly-linked protein doublecortin (DCX) by cdk5/p35 to regulate growth cone morphology and Sema3a sensitivity in developing neurons

**DOI:** 10.1101/695155

**Authors:** Christopher J. Bott, Lloyd P. McMahon, Jason M. Keil, Chan Choo Yap, Kenneth Y. Kwan, Bettina Winckler

**Affiliations:** Department of Cell Biology, University of Virginia, 1340 Jefferson Park Avenue, Pinn Hall Room 3226, Charlottesville, VA 22908, USA.; Department of Human Genetics, University of Michigan, Ann Arbor, MI 48109

## Abstract

Nestin, an intermediate filament protein widely used as a marker of neural progenitors, was recently found to be expressed transiently in developing cortical neurons in culture and in developing mouse cortex. In young cortical cultures, nestin regulates axonal growth cone morphology. In addition, nestin, which is known to bind the neuronal cdk5/p35 kinase, affects responses to axon guidance cues upstream of cdk5, specifically, to Sema3a. Changes in growth cone morphology require rearrangements of cytoskeletal networks, and changes in microtubules and actin filaments are well studied. In contrast, the roles of intermediate filament proteins in this process are poorly understood, even in cultured neurons. Here, we investigate the molecular mechanism by which nestin affects growth cone morphology and Sema3a sensitivity. We find that nestin selectively facilitates the phosphorylation of the lissencephaly-linked protein doublecortin (DCX) by cdk5/p35, but the phosphorylation of other cdk5 substrates is not affected by nestin. We uncover that this substrate selectivity is based on the ability of nestin to interact with DCX, but not with other cdk5 substrates. Nestin thus creates a selective scaffold for DCX with activated cdk5/p35. Lastly, we use cortical cultures derived from DCX knockout mice to show that the effects of nestin on growth cone morphology and on Sema3a sensitivity are DCX-dependent, thus suggesting a functional role for the DCX-nestin complex in neurons. We propose that nestin changes growth cone behavior by regulating the intracellular kinase signaling environment in developing neurons. The sex of animal subjects is unknown.

**Significance Statement:** Nestin, an intermediate filament protein highly expressed in neural progenitors, was recently identified in developing neurons where it regulates growth cone morphology and responsiveness to the guidance cue Sema3a. Changes in growth cone morphology require rearrangements of cytoskeletal networks, but the roles of intermediate filaments in this process are poorly understood. We now report that nestin selectively facilitates phosphorylation of the lissencephaly-linked doublecortin (DCX) by cdk5/p35, but the phosphorylation of other cdk5 substrates is not affected. This substrate selectivity is based on preferential scaffolding of DCX, cdk5, and p35 by nestin. Additionally, we demonstrate a functional role for the DCX-nestin complex in neurons. We propose that nestin changes growth cone behavior by regulating intracellular kinase signaling in developing neurons.

## Introduction

Correct wiring of the brain requires the targeting of growing axons to their future target fields. The outgrowth of axons is driven by the growth cone (GC) which is an actin- and microtubule (MT)-rich motile structure at the tip of growing axons (Menon & Gupton, 2016; Miller & Suter, 2018). GCs also respond directionally to a large number of extracellular guidance cues (Petrovic & Schmucker, 2015; Yamamoto et al., 2003). Guidance cues elicit local signaling responses which result in rearrangement of the GC cytoskeleton to give directional growth (Vitriol & Zheng, 2012; Dent et al., 2011), especially via guidance cue-activated kinases and their downstream substrates (Dent et al., 2011). Phosphorylation of cytoskeleton-associated proteins downstream of guidance cues is thus an important mechanism by which incoming guidance cue information is converted it into morphological rearrangements of the GC.

Most work on the cytoskeleton of axonal GCs has focused on actin filaments and MTs (Dent & Gertler, 2003; Geraldo & Gordon-Weeks, 2009; Kapitein & Hoogenraad, 2015; Tamariz & Varela-Echavarría, 2015; Yamada et al., 1970). Distinct roles for the third main cytoskeletal filament system, the intermediate filaments (IFs), during early axon formation and GC guidance are not extensively studied (Bott and Winckler, 2020), despite early promising studies in *Xenopus* neurons (Lin & Szaro, 1995,1996; Szaro et al., 1991; Walker et al., 2001). IFs are sparsely expressed in growing axons and are composed of vimentin and α-internexin (INA) prior to expression of the other neurofilaments NF-L, NF-M, and NF-H (Benson et al., 1996; Brown, 2013; Chang & Goldman, 2004; Pennypacker & Fischer, 1991; Shaw et al., 1985; Uchida & Brown, 2004; Wang et al., 2000). Once axons reach their targets, NF-M and NF-H are greatly upregulated (Laser-Azogui et al., 2015; Nixon & Shea, 1992; Yabe et al., 2003).

The atypical IF protein nestin is highly expressed in neuronal progenitor cells. We recently found lower levels of nestin protein in post-mitotic cortical neurons, especially in the distal axon. Depletion of axonal nestin leads to increased GC size and blunts sensitivity to the cdk5-dependent axonal guidance cue Sema3a (Bott et al., 2019). Our previous study thus was the first to discover a role for nestin in neurons (Bott et al., 2019), but the molecular pathways by which nestin affects growth cones remains unknown. Nestin binds the cdk5/p35 heterodimer complex (Sahlgren et al., 2003, 2006; Yang et al., 2011), but the cdk5 substrates affected by nestin are not known. The best studied example of nestin-mediated regulation of cdk5 is at the neuromuscular junction (Yang et al., 2011), but even there the physiological substrates of cdk5 affected by nestin are still unknown. In this work, we discover a neuronal nestin-regulated cdk5 substrate, namely DCX, and show that nestin’s effect on GC morphology is DCX-dependent.

DCX is a cytoskeleton-associated protein expressed in developing neurons (Ge et al., 2006; Gleeson, et al 1999; Moslehi, et al, 2017; Sapir et al., 2000; Tanaka et al., 2004; Tsukada et al., 2005; Yap et al., 2018, 2012). Mutations in human DCX cause lissencephaly, a disease characterized by defects in neuronal migration and axon outgrowth that results in profound cortical malformations (Bahi-Buisson et al., 2013). Patients exhibit intellectual disability and intractable epilepsy and often die in childhood. *In vitro*, DCX promotes MT assembly, and DCX-bound MTs are less dynamic (Moores et al., 2004). Multiple kinases regulate DCX phosphorylation (Schaar, 2004; Tanaka et al., 2004), including cdk5 (Nishimura, et al, 2014; Xie et al., 2006). Phosphorylation of DCX by cdk5 decreases the affinity of DCX for MTs and thus leads to decreased MT stability and increased MT dynamics (Bielas et al., 2007; Tanaka et al., 2004).

In this work, we identify DCX as a novel nestin-associated protein and demonstrate that cdk5 potently phosphorylates nestin-scaffolded DCX. In addition, we show that nestin’s effects on growth cone morphology and Sema3a sensitivity are DCX-dependent, thus demonstrating a functional role for the DCX-nestin complex in neurons.

## Materials and Methods

### 293 Cell culture and transfection

HEK293 cells were maintained in DMEM+10% fetal bovine serum, and all transfections were conducted using Lipofectamine 2000 (Invitrogen) according to the manufacturer’s protocol. Phosphorylation assays and IPs were performed on low passage HEK 293s in 6cm plates. Cells in low serum (4%) were transfected for 24-26hrs, and serum starved for 10-14 hours prior to lysis, to reduce background activity of mitotic kinases. Total transfection time 34 to 40 hours.

### Cos-7 cell culture and transfection

Cos-7 cells were maintained in DMEM + 10% fetal bovine serum and all transfections were conducted using Lipofectamine 2000 (Invitrogen) according to the manufacturer’s protocol. Cells were plated on, ethanol soaked, washed, round glass cell gro Fisherbrand coverslip in each 12 well plate well, at a density of 40k per well/coverslip. 2µg total of plasmid was used per transfection, and were carried out with equivalent amounts of GFP or GFP tagged construct, and empty vector myc or myc-tagged construct. The empty vector myc control is for a c-terminal myc tagged protein, thus has no start codon and is not expected to make any myc protein. Cells in low serum (4%) were transfected for 24-26hrs, and serum starved for 10-14 hours prior to lysis, to reduce background activity of mitotic kinases. Total transfection time was 34 to 40 hours.

### Western Blot

HEK 293T cells were washed with ice cold PBS, and lysed directly on the plate (6cm) into lysis buffer (500 µl); 50mM Tris HCl, 150mM NaCl, 1% NP-40, 1mM EDTA, 1mM DTT, 1mM PMSF, 1X HALT protease/phosphatase inhibitor cocktail. Extra NaF and NaOV was added in the lysis buffer for phosphorylation experiments in addition to that already in the cocktail. After incubating lysates on a rotisserie at 4° C for 30 minutes, lysate were cleared by centrifugation at 21k x g for 15 min at 4° C. We note that SPN is not well solubilized under these lysis condition, and we find that most of the SPN protein is in the pelleted material (not shown). Supernatants were diluted to 1x with 5X Laemmli buffer for western blot. Neuron cultures for Sema3a treatments where lysed directly in the 12 well plate well into 80-100 µl 1.5x laemelli sample buffer for western blot. Samples were vortexed, boiled, and spun at 18k x g for 10 minutes prior to loading. For neurons, the protein equivalent to 100-150k neurons was loaded per lane. Equivalent amounts of cells within each experiment were run on a 4-20% polyacrylamide gel. After transfer onto nitrocellulose with the BioRad Transblot Turbo, membrane was blocked with Licor TBS blocking buffer for 1 hour at room temp. Primary and secondary antibodies were diluted in 50% blocking buffer, 50% TBS, final 0.1% Tween-20. Primary antibodies were incubated overnight at 4° C, and after washing, followed by near-infrared secondary antibodies for 1 hour at room temperature. Blots were imaged with the dual color Licor Odyssey CLx imager, so each phospho-probe could be directly compared to the total protein in the same band. Densitometry of western blots was performed with Licor Image Studio software, the same software used to acquire the fluorescent blot images. The phospho antibody signal was divided by the total protein level (GFP tag or non-phospho specific antibody), and normalized to control. Positions of molecular weight ladders (in KD) are indicated next to the blot when there was room, otherwise the value of the approximate position in KD is shown.

### Immunoprecipitation

Cell lysates were prepared as though for western blotting-500ul of lysis buffer per 6cm dish of confluent cells. After the 21k x g spin, cells were precleared with 50µl of Thermo fisher control agarose for 4-6 hours. All incubations and washes were performed on the rotisserie at 4°C. For IP’s for myc-tagged proteins, 12µl Bethyl goat anti-myc pre-conjugated beads were added to the pre-cleared lysate overnight.

Lambda phosphatase dephosphorylation experiments were prepared from transfected 10cm dishes of confluents cells, and lysed with 1ml of lysis buffer (0.4% NP-40) minus the phosphatase inhibitors in the GFP-DCX condition. Following clearing by centrifugation (21k x g 15min), GFP-DCX lysates were split into two Eppendorf’s, and to one, phosphatase inhibitors were added, and to the other 3200 units (8µl) of lambda (λ) phosphatase (SCBT) and a final concentration of 2mM MnCl_2_. Both conditions were incubated at 30°C for 30 minutes, and the λ phosphatase containing sample was quenched with EDTA and phosphatase inhibitors. Each sample was separately combined each with half of the nestin-myc lysate, and incubated overnight at 4°C with 12µl Bethyl goat anti-myc beads.

For IP’s for other proteins (nestin, dcx, p35), 2µg of the relevant antibody was added to the precleared lysate for 4-6 hours, followed by addition of 12µl A/G beads overnight. Experiments using protein A/G beads for IP also used them for pre-clearing instead of the control agarose. After overnight incubation, all IPs were washed 6 times in 500µl IP buffer. For GFP nano-body precipitations for phosphorylation assays, in order to minimize time and thus loss of phospho-epitopes, no pre-clearing was performed, the GFP-trap beads were immediately added to post-21k x g spin lysates for 1hr, and washed 2x. All IPs were eluted by addition of 2x sample buffer and immediate western blotting.

### Neuronal Culture

Primary cultures of cortical neurons were obtained from embryonic day 16 (E16) mouse cortex of either sex as described in Bott et al., 2019. The protocol follows institutional ACUC protocols (approved animal protocol University of Virginia #3422). Cells were plated on acid-washed poly-L-lysine coated coverslips and incubated with Neurobasal medium with 10% fetal bovine serum and Penn/Strep. After 8 hrs, the cells were transferred into serum and antibiotic-free neurobasal medium supplemented with B27 (Invitrogen) and glutamax and cultured for a total of the indicated time periods *in vitro* (DIV). When WT vs DCX null neurons were compared within an experiment, mice were dissected and cultures prepared at the same time for consistency of comparison. For imaging purposes, cells were plated at ∼100,000 cells per 15mm coverslip or at 130,000/coverslip in electroporation experiments. Cortical neurons for culture were harvested at E16, just prior to gliogenesis. Thus non-neuronal glia are rare in these cultures, but occasional nestin/Sox2 positive, GFAP negative persisting neural stem cells were found. Cells were fixed at 24 hours (for non-transfection experiments) or 42-48 hours for overexpression experiments.

For co-IP experiments in neurons, cells were prepared similarly as described above. Six million untransfected cells were plated per 6 cm dish. At 24hours, the cells in the 6cm dish were lysed and subjected to DCX immunoprecipitation as described above. Cells for Sema3a treatments and western blot, cells were prepared similarly as described above. Untransfected cells were plated at a density of 1 million cells per 12 well plate well. After 24 hrs, cells were treated with mouse Sema3a (R&D systems) diluted in growth media for 5, 15, or 30 minutes at 1nM as indicated. All conditions were subjected to 25nM calyculinA treatment 3 minutes prior to lysis to inhibit relevant phosphatase activity prior to lysis. After treatments, media was removed thoroughly and cells were quickly lysed directly into 1.5x laemmli sample buffer and subjected to western blotting as described above. In some experiments as indicated, cells were pre-treated with 10µM Roscovitine (Cayman Chemical) for 30 minutes prior to Sema3a treatment.

### Neuron nucleofection

After cells were dissociated from E16 mouse cortex, cells were electroporated by Amaxa 4d nucleofection, according to manufacturer’s protocols. 800,000 cells were electroporated in the small volume (20µl) cuvettes using P3 primary cell solution and CL-133 program. 0.1µg of GFP plasmid (pCMV-GFP-Clonetech) as a transfection marker, and 0.15µg of relevant myc-tagged construct was used in each nucleofection condition. The cells were then all plated in a 6cm dish containing 6 poly-L-lysine coated coverslips.

### Immunofluorescence

Cells were fixed in prewarmed 4% Paraformaldehyde-PHEM-Sucrose (PPS: 60mM PIPES, 25mM HEPES, 10mM EGTA, 2mM MgCl2 (PHEM), 0.12M sucrose, 4% paraformaldehyde pH 7.4) for preservation of the cytoskeleton and cellular morphology and fixed at room temp for 16 min. Thorough permeabilization was required for nestin/intermediate filament visualization in neurons. Coverslips were permeabilized with 0.25% Triton-X 100 in 1% BSA/PBS for 20 min and then blocked for 30 min in 10% BSA, 5% normal donkey serum (NDS) in PBS all at room temperature. Primary antibodies were diluted in 1% BSA in PBS, and secondary antibodies in 1% BSA, 0.5% normal donkey serum in PBS. Primary antibody were incubated overnight at 4° C, and secondary antibodies for 90 min at room temperature. Appropriate species specific Alexa-350, -488, -568, or -647 labeled secondary antibodies raised in Donkey (Invitrogen and Jackson Immuno Research) were used for fluorescent labeling. Phalloidin-568 (Invitrogen) at 1:150 was used to visualize F-actin. Coverslips were mounted with ProLong Gold (Invitrogen). Widefield images for quantification were captured with a Zeiss Z1-Observer with a 40x objective. Images were captured with the Axiocam503 camera using Zen software (Zeiss) and processed identically within experiments. Confocal images of the whole neurons were captured with inverted Zeiss LSM880 confocal microscope using a 63X objective. Enhanced resolution images of growth cones were captured using a Nikon TiE with A1 confocal microscope with a 100x objective and NA 1.49. Images were deconvolved using Landweber for 10 iterations. No non-linear image adjustments were performed.

### *In utero* electroporation with CRISPR

The IUE CRISPR/Cas9 strategy is as follows: A gRNA sequence targeting the c-terminus of the Actb coding sequence (5’-AGTCCGCCTAGAAGCACTTG) was cloned into eSpCas9Opt1.1, expressing enhanced-specificity Cas9 and utilizing an optimized gRNA scaffold. To construct Actb_3xHA, tandem HA tag sequences were cloned between two Actb gRNA recognition sequences in the same orientation. Timed-pregnant dams were anesthetized with Ketamine (100 mg/kg) and Xylazine (10 mg/kg) via intraperitoneal injection, and given carprofen (5 mg/kg) via subcutaneous injection for analgesia. Uterine horns of an E13.5 pregnant CD-1 dam were exposed by midline incision. Lateral ventricles of embryos were injected with plasmid mix, composed of eSpCas9opt1.1-Actb_gRNA (1 μg/μl), Actb_3xHA (1.5 μg/μl), and 0.1% Fast Green FCF to visualize solution during intraventricular injection. Successful genome editing and repair in a subset of transfected cells yielded an HA-tagged Actb locus and led to sparse expression of ACTB-3xHA in individual cells. Electroporation paddles were positioned on opposing sides of the ventricle and 4 - 5 pulses of 27 volts (45 milliseconds each) were delivered, with 950 milliseconds between pulses, delivered using a BTX Harvard Apparatus ECM 830 power supply. Brains were isolated 3 days after electroporation, at E16.5, and fixed overnight in 4% PFA. These experiments were carried out in compliance with ethical regulations for animal research. The study protocol was reviewed and approved by the University of Michigan Institutional Animal Care & Use Committee.

### Histology

Fixed *in utero* electroporated E16.5 mouse brains were then washed with PBS, embedded in agarose and vibratome sectioned into 100 um sections. Sections were stained as floating sections-permeabilized and blocked with 3% normal donkey serum + 0.3% Triton-X 100 in PBS for 1 hour at room temperature. Antibodies were diluted in the same permeabilization buffer with indicated primary and secondary antibodies sequentially overnight at 4°C. Sections were thoroughly washed over a period of 10hrs at room temperature with 0.5% BSA and 0.1% Triton-X 100 in PBS after primary and secondary antibody incubations. Sections were mounted on slides with Prolong Gold using 22X22mm 1.5 fisher brand coverslips. Confocal and Airyscan imaging of cryosections was carried out on an inverted Zeiss LSM880 confocal microscope (available in the UVA microscopy core facility) using a 63X objective.

### Molecular biology

Plasmids

GFP-DCX (Yap et al., 2016)

Negative control substrates-GFP tagged:

DCLK1 (Shin E., et al., 2013)
CRMP2A (Balastik M., et al., 2015)
FAK (addgene)
Drebrin-YFP (addgene)
Tau-YFP (George Bloom)

P35 (addgene)

HA-CDK5 (addgene)

HA-CDK5-DN (D145N) (addgene)

Myc-SPN Lawrence Brass (UPenn)(Yap et al., 2016)

INA-myc (Sino biological)

Nestin-myc (Bott C., et al, 2019 via original cDNA Bruce Lahn) PCR was used to generate EcoR1 and HindII sites on the 5’ or 3” end respectively of full length mouse nestin. Both the insert and vector were treated with EcoR1 and HindII, the cut insert was ligated into pCDNA3.1 Myc-His(+) B.

Myc EV 3.1 B (stratagene)

GFP (Clonetech)

CMV-HA EV-(addgene)

#### New constructs made for this study

Nestin mutants:

<u>Nestin-myc T316A</u>-We used to traditional primer mutagenesis for this plasmid derived from the pcDNA 3.1 B Nestin(wt)-myc plasmid. Only one nucleotide change was needed to mutate T(946ACA) to A(946GCA). Primer: 5’-gaacttcttccaggtgcctgcaagcgagagttc-3’

<u>Nestin-myc T316D</u>-Could not be obtained via traditional mutagenesis and was made, instead via gene synthesis from genscript of a large >1kb piece of nestin that included the three nucleotide changes-T(946ACA) to D(946GAT). The 5’ end had a HINDIII site for incorporation into the MCS of pcDNA 3.1 B nestin(wt)-myc plasmid. An ECOR1 site present in WT nestin was used for the 3” end.

#### Image analysis

All image analysis was performed on ImageJ blinded to experimental conditions. Transfected neurons were identified by having a prominent GFP fluorescence. Stage3 neurons were identified (according to Banker staging) by having one neurite (the future axon) at least 2x longer than the next longest neurite. The distal portion of the future axon was analyzed for morphological measurements as the “axon growth cone”.

Images were analyzed as in Bott et al. 2019.

*Filopodial protrusions* were counted using the phalloidin channel, and any protrusions from the growth cone were counted.

*Axon length* was the length of the axon from the cell body to the growth cone tip as in Bott et al. 2019.

*Growth cone area* were assessed by measuring traced outlines the growth cone not counting the filopodial protrusions, and extending down proximally to the point where the splayed growth cone microtubules consolidate into bundles in axon shaft as in Bott et al. 2019.

*For transfected Cos7 cells*: Pearson correlation coefficients were determined using the Coloc module in Imaris 9.5.1. Regions of interest were defined as the entire area within a transfected cell expressing the two markers of interest, then the Pearson value was determined within that ROI (termed “Pearson’s coefficient in the ROI volume” in Imaris).

### Statistical analysis

All data was statistically analyzed using Prism software. Datasets were first evaluated for normality by using the Shapiro-Wilk normality test. This result was used to decide if parametric or non-parametric tests were to be used. When more than one comparison was made, the corresponding ANOVA test was used. When only two conditions were compared, we used a t-test. Figure legends specify details-for each set of experiments we made an effort to show precise p-values when shown. p-values reaching statistical significance of at least <0.05 are in **bold**.

### Microscope image acquisition

Widefield images for quantification were captured with a Zeiss Z1-Observer with a 40x objective. Images were captured with the Axiocam503 camera using Zen software (Zeiss) and processed identically within experiments.

Confocal images of the whole neurons or brain sections were captured with inverted Zeiss LSM880 confocal microscope using a 63X objective with Zen software.

Airyscan was used for some brain section imaging.

Enhanced resolution images of growth cones were captured using a Nikon TiE with A1 confocal microscope with a 100x objective and NA 1.49. Images were deconvolved using Landweber for 10 iterations. No non-linear image adjustments were performed.

### Reagents

#### Antibodies

Mouse anti-Nestin 2Q178 1:200 IF, 1:500 IHC, 1:500 WB **Santa Cruz Biotechnology Cat# sc-58813 RRID:AB_784786**

Goat anti-Nestin R-20: Raised against peptide in the unique C-terminal tail region of nestin 1:200 IHC **Santa Cruz Biotechnology Cat# sc-21249 RRID:AB_2267112**

Rabbit anti-DCX-1:1200 IF, IHC, 1:4000 WB **Abcam Cat# ab18723 RRID:AB_732011**

Mouse anti-Β III Tubulin Tuj1-1:800 IF A generous gift from the Deppman Lab YVA **A.**

**Frankfurter, University of Virginia; Department of Biology Cat# TuJ1 (beta III tubulin) RRID:AB_2315517**

Chicken anti-βIII tubulin-1:200 IF **Aves Labs Cat# TUJ RRID:AB_2313564**

Rat anti α-tubulin-1:2000 WB **Santa Cruz Biotechnology Cat# sc-53030**

**RRID:AB_2272440**

Rabbit anti-Vimentin-1:100 IF, 1:100 IHC, 1:1000 WB **Bioss Inc Cat# bs-0756R RRID:AB_10855343**

Rabbit anti-Vimentin 1:1000 IF **Abcam Cat# ab92547, RRID:AB_10562134**

Mouse anti-Myc tag 9E10: 1:1000 WB, 1:200 IF **Santa Cruz Biotechnology Cat# sc-40, RRID:AB_627268**

Rat anti-Myc tag 9E1-1:1000 WB ChromoTek Cat# 9e1-100, RRID:AB_2631398

**Rat anti-HA 1:2000 WB ChromoTek Cat# 7c9, RRID:AB_2631399**

Rabbit anti-GFP D5.1 1:5000 WB **Cell Signaling Technology Cat# 2956, RRID:AB_1196615**

Mouse anti-GFP N86/8 1:5000 WB **UC Davis/NIH NeuroMab Facility Cat# 73-131, RRID:AB_10671444**

Rabbit anti-CDK substrate multiMab 1:1000 WB **Cell Signaling Technology Cat# 9477, RRID:AB_2714143**

Rabbit anti-pS297 DCX 1:1000 WB **Cell Signaling Technology Cat# 4605, RRID:AB_823682**

Rabbit anti-pS334 DCX 1:2000 WB **Cell Signaling Technology Cat# 3453, RRID:AB_2088485**

Mouse anti-CDK5 1:2000 WB **1H3 Cell Signaling Technology Cat# 12134, RRID:AB_2797826**

Rabbit anti-p35/p25 1:2000 WB **C64B10 Cell Signaling Technology Cat# 2680, RRID:AB_1078214**

Mouse anti-Spinophilin (Neurabin II) 1:2000 WB **Santa Cruz Biotechnology Cat# sc-373974, RRID:AB_10918794**

Mouse anti-Drebrin 1:2000 WB **Santa Cruz Biotechnology Cat# sc-374269, RRID:AB_10990108**

Mouse anti-Phf1 tau-supernatant 1:50 WB **P. Davies Albert Einstein College of Medicine; New York; USA Cat# PHF1, RRID:AB_2315150. A gift from George Bloom.**

Rabbit anti-pS522 Crmp2a 1:1000 WB **Thermo Fisher Scientific Cat# PA5-37550, RRID:AB_255415**

Mouse anti-pS732 FAK 1:1000 WB **Santa Cruz Biotechnology Cat# sc-81493, RRID:AB_1125825**

Rabbit anti-pY397 FAK 1:1000 WB **Bioss Cat# bs-1642R, RRID:AB_10856939**

Gt anti-myc agarose 1:50 IP **Bethyl Cat# S190-104, RRID:AB_66618**

GFP trap agarose 1:100 IP **ChromoTek Cat# gta-20, RRID:AB_2631357**

GFP booster nanobody-1:600 IF **ChromoTek Cat# gba488-100 RRID:AB_2631434**

Phalloidin 568 1:150 Invitrogen

#### Cell lines

Hek293 cell lines: **ATCC Cat# PTA-4488, RRID:CVCL_0045**

Cos7 cell lines: **ATCC Cat# CRL-1651, RRID:CVCL_0224**

## Results

### Nestin selectively augments cdk5-phosphorylation of DCX, but not of other cdk5 substrates

Our recent work (Bott et al., 2019) discovered an unexpected role for nestin in axonal growth cones: downregulation of nestin increases GC size and decreases Sema3a sensitivity. In other cell types, nestin has been shown to affect the activity of cdk5. In neuromuscular junction (NMJ) development, for example, nestin deletion phenocopies cdk5 deletion or treatment with the cdk5 inhibitor roscovitine by altering levels and localization of cdk5 activity (Lin et al., 2005; Yang et al., 2011). Therefore, we hypothesized that nestin acted via cdk5 in early neurons. Cdk5 is important for many processes in neurite outgrowth, including responses to guidance cues (Dhavan & Tsai, 2001; Kawauchi, 2014). Knockout mouse models that lack cdk5 produce a cortex with disorganized layering, similar to what is seen in lissencephaly (Tanaka et al., 2004), and result in perinatal lethality (Oshima et al., 1996; Hirasawa et al., 2004). Other studies have shown that balancing cdk5 activity is important, as increasing cdk5 function proves also disruptive (Cheung & Ip, 2012). Since nestin binds the cdk5 activator p35 (Sahlgren et al., 2003, 2006; Yang et al., 2011), and cdk5 inhibition abrogates the effect of nestin-associated Sema3a-mediated GC collapse in neurons (Bott et al., 2019), we reasoned that nestin might change the phosphorylation of one or more cdk5 substrates to mediate the nestin-dependent changes in axonal GCs. We thus tested whether the phosphorylation of several known neuronal cdk5 substrates was affected by the expression of nestin.

To this end, we utilized an “in-cell” phosphorylation assay in transfected HEK293 cells (Fig. 1A-E). HEK293 cells endogenously express cdk5, but not the cdk5 activator p35 (see cdk5 and p35 immunoblot in Fig. 2A). They thus have minimal cdk5 activity under basal conditions. We focused on Focal Adhesion Kinase (FAK), Collapsin Response Mediator Protein 2 (Crmp2), tau, and DCX since they are known to be phosphorylated in neurons downstream of Sema3a signaling and phospho-specific antibodies are commercially available (Fig. 1A-E). Transfection of HEK293 cells with p35 to activate cdk5 led to increases in phosphorylation of GFP-tagged FAK, Crmp2, or tau, as expected (Fig. 1A,C,D). Surprisingly, p35 expression was not sufficient to increase cdk5-mediated phosphorylation of DCX in this context (Fig. 1E). When nestin was expressed in addition to p35; FAK, Crmp2, or tau showed no further increase in phosphorylation. However, DCX phosphorylation became significantly increased when nestin was co-expressed with p35. A myc-empty vector (myc-EV) was used as a negative control. For quantification of all conditions, we used ratios of the phospho-protein signal to total expression of the GFP-tagged substrate protein (Fig. 1A’-E’). As an additional control, we utilized an antibody to a phospho-tyrosine residue on FAK. As expected, pY397-FAK was not affected in any condition (Fig. 1B, B’). We note that there was basal phosphorylation for some of these substrates even in the absence of overexpressed p35, which is likely due to activity of other kinases in HEK293 cells. We thus discovered a surprising substrate selectivity for nestin-enhanced phosphorylation by p35/cdk5. This finding suggested the hypothesis that DCX could be a downstream effector for the nestin-dependent effects seen in the growth cones of young neurons.

**Figure 1:**
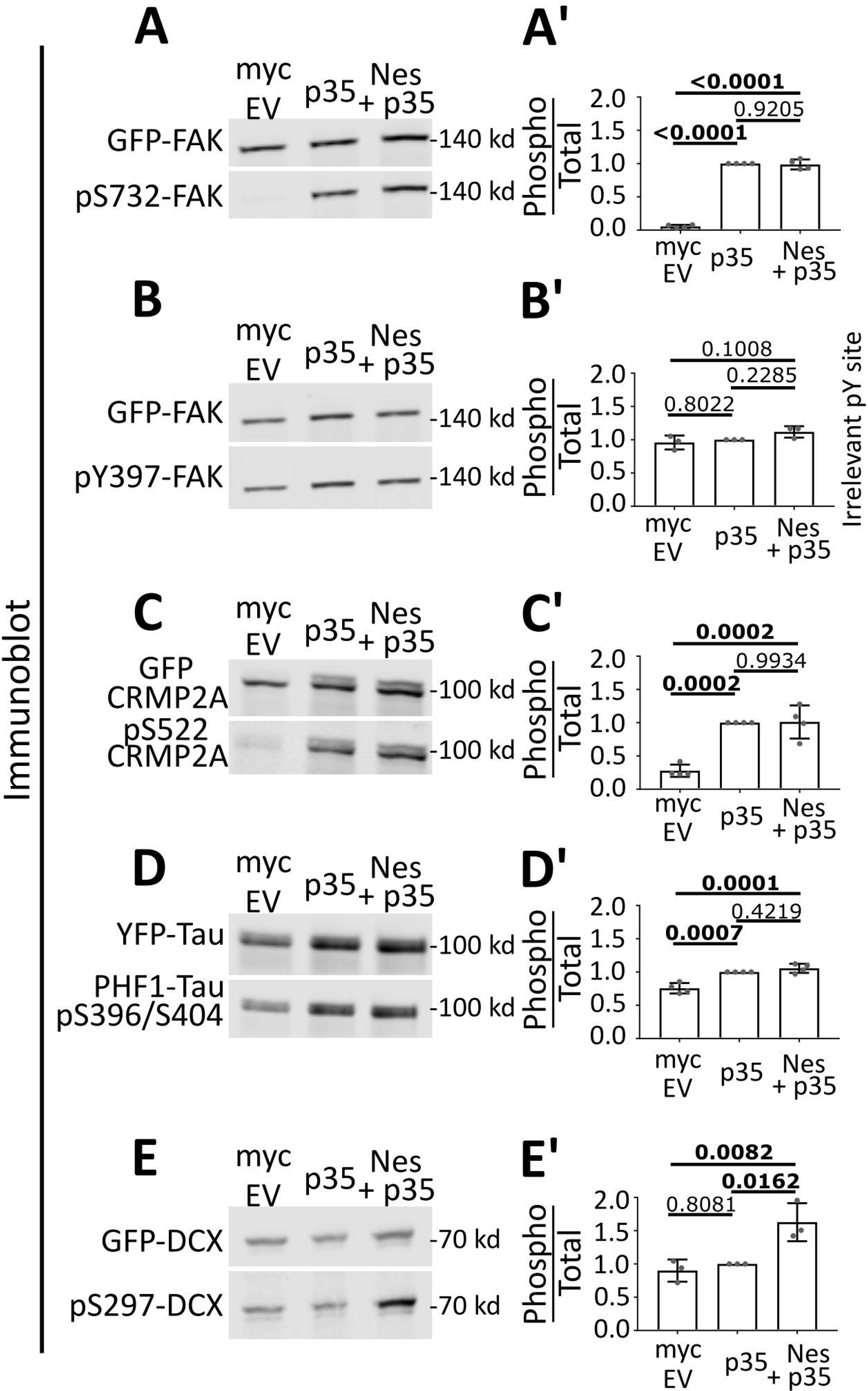
Nestin selectively augments CDK5-phosphorylation of DCX, but not other cdk5 substrates. A-E. Comparison of cdk5-mediated phosphorylation in HEK293 cells of several cdk5 substrates in the indicated transfection conditions: Myc-EV (empty vector) – left lane, Myc-EV + p35 – middle lane, Nes-myc + p35 – right lane. Cdk5-dependent phosphosites on FAK (A), Crmp2 (C), tau (D) and DCX (E) were assessed for p35- and nestin-dependence. The pY397 site on FAK (B) is a negative control because cdk5 is a Ser/Thr kinase. A’-E’. The level of phosphorylation of each substrate was quantified by densitometry analysis of the phospho-signal compared to total (GFP) and graphed to the right of each blot. Error bars are SD. The statistical test used was one-way ANOVA with Tukey’s correction. N= 3 or 4 independent experiments. *p* values of less than 0.05 are indicated in bold.

**Figure 2:**
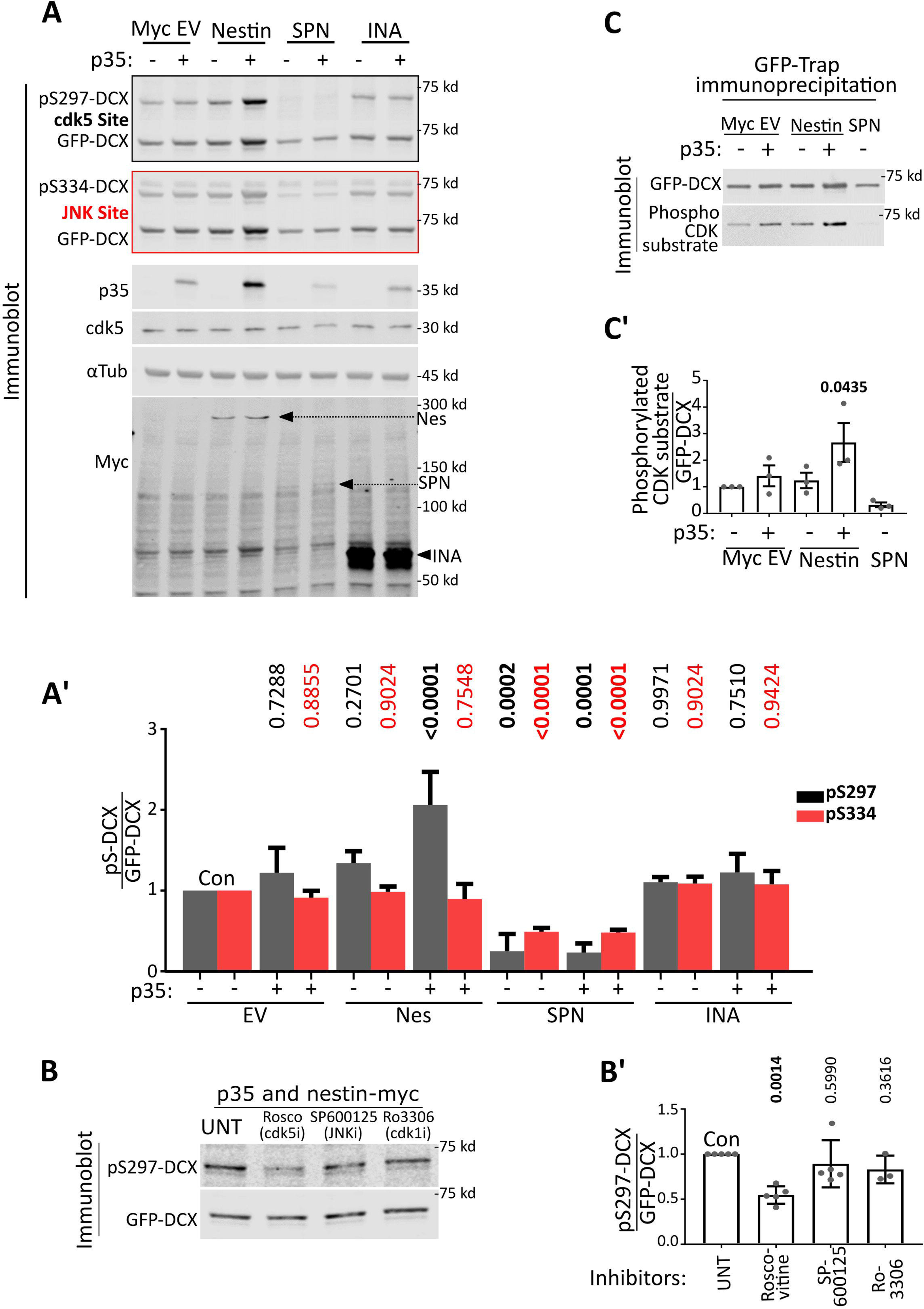
Nestin, but not α-internexin, promotes phosphorylation of DCX by cdk5. A. Myc Empty Vector (EV), Nestin-myc, INA-myc, or myc-SPN were expressed in HEK293 cells with DCX alone (-), or with both p35 and DCX (+). Cell lysates were run on two replicate western blots and both blotted against GFP for total levels of GFP-DCX. Each was blotted with the relevant phospho-specific antibodies to pS297-DCX (a cdk5 site) or pS334-DCX (a JNK site). The levels/presence of relevant proteins are shown, including ckd5 (endogenous), p35 (overexpressed), loading control), and the overexpressed myc-tagged constructs. Arrows point at the relevant myc-tagged protein bands. A’. Quantification of A by densitometry. Ratio of phospho-DCX to total DCX are plotted. Black bars represent pS297-DCX/total DCX, and red bars represent pS334-DCX/total DCX-colors correspond to the outline around the relevant blot. Error bars are SD. N = 4 to 6 independent experiments. Every condition was compared to every other condition with an ordinary one way ANOVA test with Tukey correction. P-values compared pairwise to control (CON) are shown. Additional comparisons not shown are pS297 level in Nestin + p35 condition is significantly different than p35 alone (p<0.0001) and nestin alone (p=0.0004), while pS334 is not significantly different (>0.9999 and 0.8979 respectively) for the same comparisons. *p* values of less than 0.05 are indicated in bold. The values for DCX phosphorylation are from additional independent experiments separate from those shown in Fig. 1E. B. HEK293 cells transfected simultaneously with p35, nestin-myc, and GFP-DCX were treated with inhibitors (i) of cdk5 (roscovitine), JNK (SP600125), cdk1 (Ro2206), or vehicle (UNT) for 6h before lysis. Only roscovitine significantly decreased levels of pS297-DCX. B’. Quantification of the blots in B. Error bars shown are SD, ordinary one-way ANOVA with Dunnett’s correction was performed and the pairwise p-values are shown for each condition compared to untreated control (UNT). N=3-5 independent experiments. *p* values of less than 0.05 are indicated in bold. C. HEK293 cells were transfected similarly to Figure 2A and total cdk5-mediated phosphorylation of GFP-DCX was assessed using an antibody to a generic K/H SP cdk5 phosphorylation motif to detect. Because many proteins in HEK293 cell lysates can be recognized with this antibody, cell lysates were subjected to GFP-trap incubation and precipitation to isolate GFP-tagged DCX from other reactive proteins and subsequently subjected to western blot. C’. Densitometry quantification of C where levels of signal to the phospho-reactive antibody is ratio-ed to the level of total DCX. Error bars are SD and a one way ANOVA with Dunnett’s correction was used to compare each condition to the control cells transfected with GFP-DCX only. N= 3 independent experiments. *p* values of less than 0.05 are indicated in bold.

### Nestin, but not α-internexin, promotes phosphorylation of DCX by cdk5

In order to more fully characterize the effect of nestin on DCX phosphorylation, we repeated the phosphorylation assay of DCX with additional controls. As shown above, we saw a significant increase in pS297-DCX levels only when p35 and nestin are both co-expressed (Fig. 2A,A’, grey bars). In contrast to the phosphorylation of S297, the phosphorylation of S334 by JNK was not changed by co-expression of nestin (Fig. 2A,A’, red bars). In order to test specificity of the phospho-specific antibodies, we expressed spinophilin (SPN) (Fig. 2 A,A’) which is a known PP1 phosphatase scaffold and promotes dephosphorylation of both the cdk5 and JNK sites of DCX (Bielas et al., 2007; Shmueli et al., 2006; Tsukada et al., 2006). The levels of both pS297-DCX and pS334-DCX were greatly diminished by SPN co-expression. We note that nestin co-expression with p35 consistently resulted in an increase in total DCX levels. Again, we used the ratio of the phospho-DCX signal to total expression of GFP-DCX to normalize for input levels of DCX.

We next carried out several controls to ascertain that the increased phosphorylation of DCX was attributable to cdk5 activity and not to other endogenous kinases present in HEK293 cells. The cdk5 inhibitor roscovitine reduced pS297-DCX phosphorylation, but neither SP600125 (a JNK inhibitor) nor Ro3306 (a cdk1/cdc2 inhibitor with minimal cross-reactivity with cdk5) changed the pS297-DCX levels (Fig. 2B,B’), supporting previous literature that S297 is a cdk5 site on DCX (Tanaka et al., 2004). Multiple residues in addition to S297 are phosphorylated by cdk5 on DCX, but specific antibodies to detect these phosphorylation events are not available. We obtained another antibody, which recognizes the general “K/H pSP” phospho-epitope consensus site for CDKs to confirm our results with the pS297-DCX antibody. GFP-DCX was precipitated from lysates with GFP-trap agarose beads to separate DCX from other reactive endogenous cdk substrates in this experiment. Similar to our results using the anti-pS297DCX antibody, a significant nestin-dependent increase in cdk5 phosphorylation of DCX could also be seen with this cdk phospho-substrate antibody (Fig. 2C and quantified in 2C’). Again, expression of SPN to recruit PP1 to DCX abolished the band on Western blot.

Lastly, we tested a different neuronal IF protein, α-internexin (INA) which is not known to bind to p35. INA did not promote DCX phosphorylation by p35/cdk5, arguing for a specific enhancement of DCX phosphorylation by nestin (Fig. 2A,A’).

### Nestin forms a complex specifically with DCX

Next, we asked by what mechanism nestin might selectively enhance cdk5-phosphorylation of a specific cdk5 substrate. We first tested if nestin formed a complex with any of the cdk5 substrates. We co-transfected Nestin-myc or myc-empty vector (EV-negative control) with several GFP-tagged cdk5 substrates (DCX, Doublecortin-like-Kinase 1 (DCLK1), CRMP2, Tau, Drebrin, and FAK) individually in HEK293 cells, followed by cell lysis and anti-myc tag immunoprecipitation (Fig. 3A). DCX, CRMP2, Tau, Drebrin and FAK are cdk5 substrate proteins known to be downstream of Sema3a. We used DCLK1 as an additional control since it shares sequence homology with DCX, and is a cdk5 substrate (Contreras-Vallejos et al., 2014). The efficiency of co-IP with nestin-myc was quantified for each GFP-tagged substrate by densitometry analysis as a ratio of immunoprecipitated protein to the total amount of protein in the corresponding input Low levels of other substrates, such as DCLK1 and CRMP2, were also seen in the co-IPs, but not at significant levels relative to the empty vector control. Only GFP-DCX significantly co-immunoprecipitated with nestin-myc (Fig. 3A, A’).

**Figure 3:**
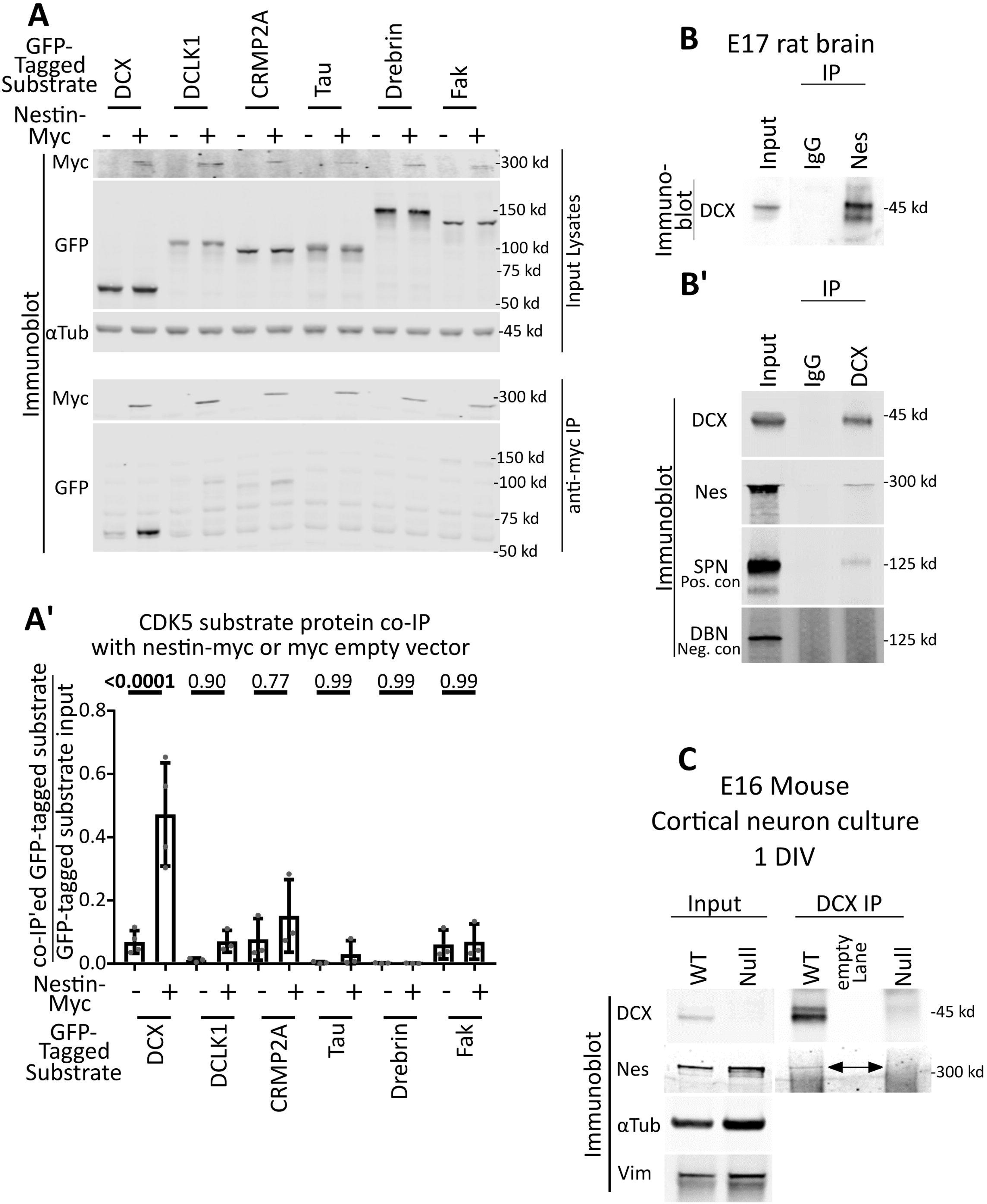
Nestin forms a complex specifically with DCX. A. Candidate screen for cdk5 substrates that interact with nestin by co-immunoprecipitation (co-IP). Six different GFP-tagged cdk5 substrates were expressed with or without nestin-myc in HEK293 cells and subjected to immunoprecipitation with anti-myc antibody. An empty vector was used as a negative control. Top panel: Input αTubulin is used as a loading control. Bottom panel: Anti-myc IPs are probed against myc or GFP. Only GFP-DCX co-IPs with nestin-myc. A’. Densitometry analysis of each co-IP’ed substrate relative to its input. Statistical comparisons (ordinary one-way ANOVA with Sidak’s correction) were performed between the empty vector myc IP compared to nestin-myc IP. Errors bars are SD, n = at least 3 for each condition. P-values are indicated above the graph. *p* values of less than 0.05 are indicated in bold. B -B’. IP of endogenous nestin and DCX complexes from brain. B. DCX co-IP with anti-nestin antibody from E17 rat brain compared to a control IgG. B’. Nestin co-IP with anti-DCX antibody from E17 rat brain compared to a control IgG. As a positive control, DCX IPs were probed against SPN. Drebrin, serves as a negative control. C. anti-DCX immunoprecipitation from 1DIV cortical E16 mouse neurons results in co-IP of nestin from WT neurons, but not from DCX knockout (null) neurons, demonstrating specificity of the IP. The double-headed arrow points at the band corresponding to nestin. No anti-DCX immunoreactivity is seen in the western blots of DCX null neurons compared to WT, also demonstrating specificity of the antibody. immunoblots are shown as loading controls.

To determine if the nestin-DCX complex occurred endogenously in the relevant tissue, lysates from embryonic rodent brain were subjected to IP with either anti-nestin (Fig. 3B) or anti-DCX (Fig. 3B’) antibodies and compared to non-immune IgG. An endogenous complex of nestin and DCX was recovered for both IPs. IP with anti-nestin antibody resulted in efficient co-IP of DCX. In contrast, relatively low amounts of nestin were recovered in the IP with an anti-DCX antibody, but when compared to the known DCX interacting protein SPN (a positive control), similar levels of co-IP efficiency were detected. This low efficiency is likely due to the fact that DCX has a multitude of binding interactions which might be mutually exclusive. The non-interacting protein drebrin was not detectable in the co-IP, serving as a negative control. In agreement with these findings, we also found nestin by mass spectrometry in DCX IPs of embryonic rat brain (Yap CC, unpublished data).

Since brain contains multiple cell types, we repeated the IP with lysates from E16 cortical neurons cultured for one day *in vitro* (DIV1). Indeed, nestin co-IP’ed with DCX (Fig. 3C) in DIV1 cortical cultures. As a negative control, we carried out IPs from neuronal cultures prepared from DCX KO E16 cortex. The DCX KO neurons expressed similar levels of nestin as WT cultures, but no nestin was detected in the DCX IP (Fig. 3C). We thus discovered a novel endogenous complex of DCX and nestin in immature cortical neurons.

Since DCX is MT-associated and nestin is an IF, we wondered where a complex of nestin with DCX would localize in cells. In order to test if DCX and nestin affected their respective cellular localizations, we transfected COS-7 cells which are large, well-spread cells which have spatially resolvable MT and vimentin IF filament systems. As shown previously by us and others, GFP-DCX decorates MTs (Ge et al., 2006; Gleeson et al., 1999; Moslehi et al., 2017; Sapir et al., 2000; Tsukada et al., 2005; Yap et al., 2018, 2012), but it is not co-localized with vimentin (Fig. 4A). Conversely, Nes-myc co-localized with endogenous vimentin in the absence of DCX (Fig. 4B). When DCX was co-expressed, nestin became enriched on DCX-decorated MT bundles (Fig. 4C). When a DCX-related MAP, GFP-DCLK1, was expressed instead of DCX, nestin remained associated with vimentin IFs (Fig. 4D). Other IFs also expressed in early neurons, such as INA, did not enrich with DCX-MT bundles (Fig. 4E). Pearson’s coefficient of co-localization was quantified for double transfected Cos7 cells, demonstrating strong co-localization between DCX and nestin, and significantly less colocalization between DCLK1 and nestin, or DCX and INA (Fig. 4F). Nestin can thus be recruited to DCX-decorated MTs.

**Figure 4:**
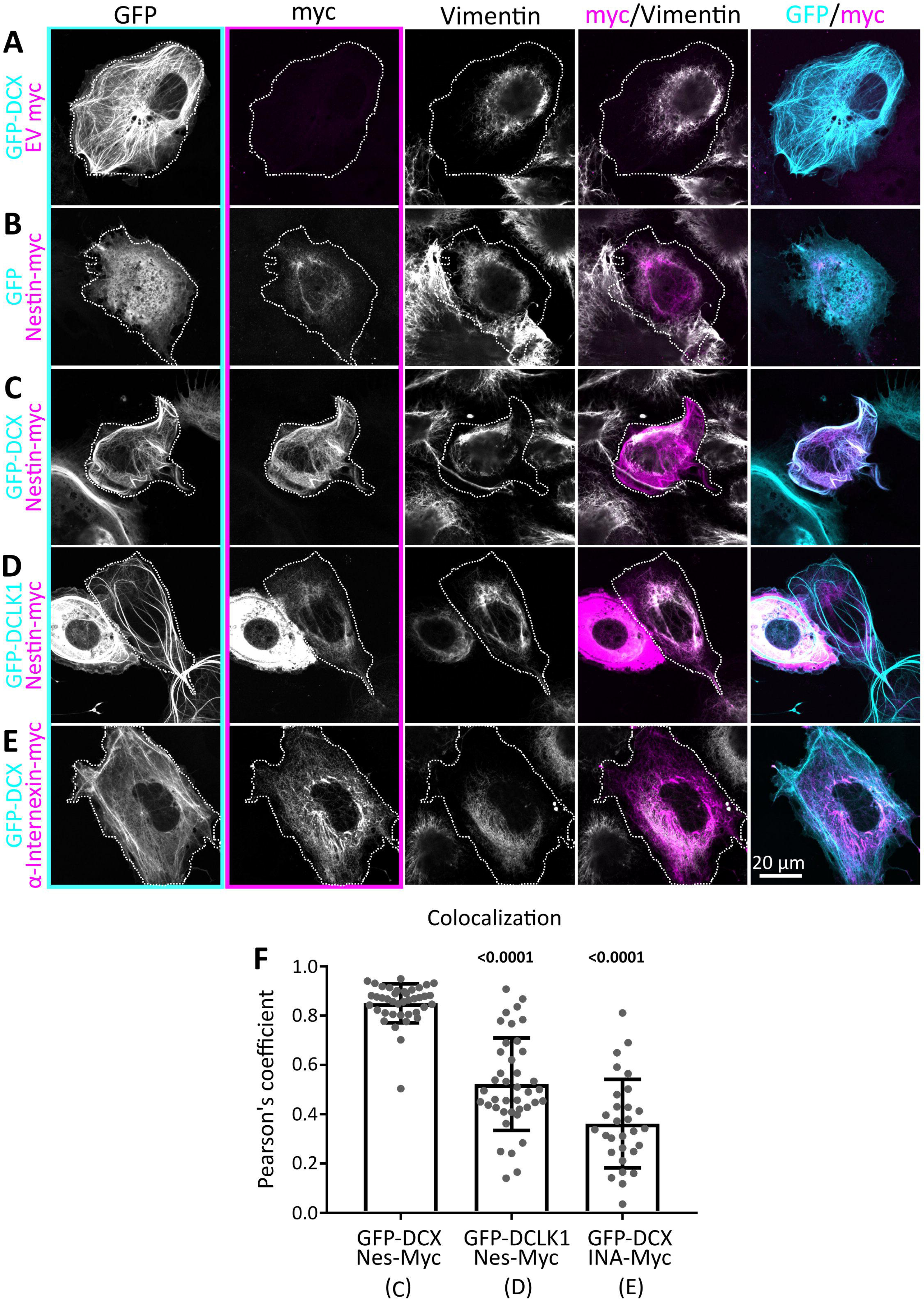
Nestin co-localizes with DCX-MT bundles. A-E: Confocal images of COS-7 cells transfected with GFP-DCX + myc EV (A), GFP + nes-myc (B), GFP-DCX + nes-myc (C), GFP-DCLK1 + nes-myc (D), and GFP-DCX+ INA-myc (E). All cells are counterstained with GFP booster nanobody, anti-myc tag antibody, and antibody against endogenous vimentin, as labeled above each panel column. The transfected cell is outlined with a dotted line. Individual black and white images are shown for each channel separately, as well as merged images of the GFP/myc and myc/vimentin channels. Panel outlines of black and white images correspond to the color used in the composite images. Nestin is closely co-localized with vimentin in the perinuclear region, except when GFP-DCX is co-expressed which enriches nestin onto DCX MT bundles. Vimentin remains in the perinuclear region. DCLK1 and internexin do not have the same relationship with Nestin or DCX, respectively. F: Pearson’s coefficient of conditions represented above in C, D, and E (as indicated below the x-axis labels. N=30 to 40 cells from 2-3 independent experiments are shown with the mean, and error bars are SD. Data was analyzed with non-parametric Kruskal-Wallis one way ANOVA test with Dunn’s correction, with each p-value shown above each condition compared to GFP-DCX with Nes-myc (1^st^ lane). Both p-values are less than 0.05 and are thus in bold. Both nestin with DCLK1, and INA with DCX have significantly less colocalization, as indicated by the smaller Pearson coefficients.

### Nestin and DCX co-localize in the distal axon of cortical neurons *in vitro* and *in vivo*

We previously described a striking localization of nestin in the distal axon of cultured E16 mouse cortical neurons at 1 DIV that is rapidly lost as neurons matured (Bott et al., 2019), and nestin mRNA has recently been found to be enriched in neurites (Kügelgen & Chekulaeva, 2020). DCX is enriched in the distal axon, but is also present in dendrites (Friocourt et al., 2003; Schaar et al., 2004). Simultaneous staining with antibodies against DCX and nestin revealed co-localization in the distal axon (Fig. 5A; arrowhead), but not in dendrites. Enhanced resolution microscopy of the distal axon showed that nestin co-distributed with DCX in the “wrist” region of the axonal growth cone (Fig. 5B) and appeared to be filamentous. Nestin staining also extended past the “wrist” into the central domain of the GC. Interestingly, we observed occasional nestin staining in some of the proximal filopodia of the growth cone (arrowhead in Fig. 5B) which were also stained for DCX.

**Figure 5:**
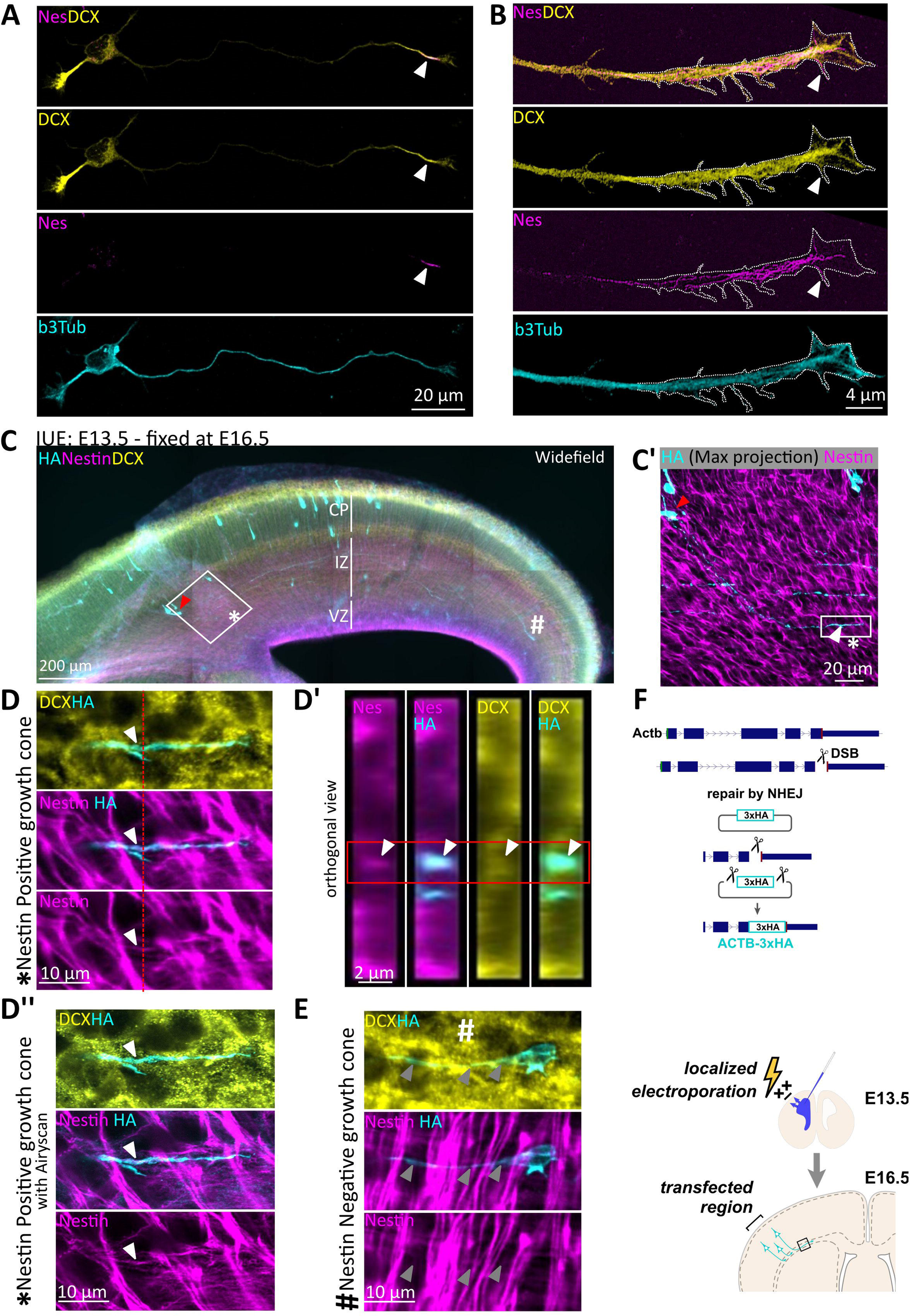
Nestin co-localizes with DCX in axons of newly born neurons *in vitro* and *in vivo*. A. Confocal images of an E16 cortical mouse stage 3 neuron after 1 DIV culture, counterstained for endogenous DCX, nestin, and β3 tubulin. DCX is enriched aty the distal region of all neurites, whereas nestin immunostaining labels only the distal region of the growing axon (arrowhead), but not of the minor neurites. B. Enhanced resolution confocal imaging of nestin in the distal region of an axon-together with DCX. Distally, some nestin filaments appear to extend into the central region of the growth cone and into some proximal filopodia (arrowhead). C. Wide field immunofluorescence image of a coronal section of E16.5 mouse cortex to illustrate the sparse labeling of individual neurons achieved with knockin of HA-tag using CRISPR and IUE (*in utero* electroporation). In all panels, red arrowhead indicates the cell body that projects the HA/Nes/DCX positive growth cone. “*” marks the nestin positive growth cone, whereas # marks the nestin negative neuron (shown in D”, E). The major cortical layers are labeled: VZ-ventricular zone, IZ-intermediate zone, CP-cortical plate. Antibody labeling is indicated in each panel. C’. Inset of square in C demonstrating HA-staining of a single stage 3 neuron with the entirety of its axon imaged in a confocal stack (maximum projection). The distal axon/growth cone is marked with a white arrowhead, and cell body with a red arrowhead. D and D’ panels are confocal images, and D’’ is an airyscan enhanced resolution image. D. High magnification zoom of the inset square of C’ of the growth cone marked with * in C and C’. A single z-plane of is shown. D’. Orthogonal projection of the region indicated in the red dashed line of D. The red box outlines the relevant HA-labeled cell, which is positive for nestin and DCX. D’’. High magnification Zeiss Airyscan confocal image equivalent of D. E. High magnification zoom of the growth cone marked with “#” in C. Arrowheads point along the length of the axon, nestin is not detected in this growth cone. F. Diagram of the experimental strategy to detect nestin and DCX in single growth cones in mouse cortex. E13.5 mouse cortex was electroporated *in utero* to sparsely introduce three HA-tags into β-actin (Actb-3xHA) by CRISPR. Brains were fixed at E16.5. The CRISPR strategy is depicted diagrammatically.

We previously showed nestin immunostaining in E16.5 mouse cortex in axon fascicles in the intermediate zone (IZ) which are interwoven among highly expressing radial glial processes (Bott et al., 2019). However, we were unable to discern single axons *in vivo*, let alone single growth cones, due to the high density of axons in the IZ. In order to visualize nestin localization in individual developing axons *in vivo*, we utilized *in utero* electroporation (IUE) to sparsely label individual neurons (Fig. 5C). Using CRISPR/Cas9 genome editing, 3xHA tags were introduced into the endogenous β-actin locus with enabled us to label the actin cytoskeleton of developing neurons including the growth cone (diagram in Fig. 5F). Three days after IUE, brains were fixed, sectioned and immunostained for the HA-tag in combination with staining for DCX and nestin (Fig. 5C). This technique introduced the HA tag into a small number of newly born neurons, which were scattered throughout the IZ and cortical plate (CP) at multiple stages of maturation. We expected to find nestin-positive axon tips in only a subset of labeled neurons (Bott et al., 2019) and therefore focused on neurons that were migrating though the IZ and had not yet reached the cortical plate (Fig. 5C, white asterisk “_*_”). These neurons are less mature and still in the early stage of axon extension (Namba et al., 2014; Sakakibara & Hatanaka, 2015) and thus most analogous to the young neurons we used (and quantified in Bott et al., 2019) in culture. One such neuron (soma marked with red arrowhead, axon tip marked with white arrowhead) is shown enlarged in Fig. 5C’. The nestin staining pattern in the IZ consisted of bright radial staining of the radial glial processes and fainter staining of axons running parallel to the ventricular surface, as we showed previously (Bott et al., 2019). The sparse labeling provided by the HA-tag knock-in strategy thus allowed us to identify individual neurons at the earliest stage of axon extension and positioning and to assess for nestin staining. A high magnification of a single section of a z-stack shows an example of nestin staining that clearly localized within the distal region of the extending axon (Fig. 5D), but did not extend to the most distal regions of the growth cone. Orthogonal projections of the z-stack were used to allow for visualization of nestin within the HA-positive axon (Fig. 5D’). Enhanced resolution Airyscan confocal imaging of the same growth cone in the same z-plane allowed for better appreciation of the axonal nestin (Fig. 5D’’). Nestin in distal axons was thus also observable *in vivo* and thus mirrors what was seen in culture (Bott et al., 2019). DCX protein localization was observed in the same nestin-positive process, but also more broadly in other axons, i.e. throughout different stages of neuron maturation, and is thus seen in more neurons than nestin. We observed nestin primarily in the smaller, more compact GCs of younger neurons whose cell bodies had not yet reached the cortical plate (5C “*” symbol). The growth cones of longer axons (presumably belonging to more mature neurons) did not have detectable nestin enrichment and were larger in size (Fig. 5E, 5C “#” symbol). Even though we were able to observe examples of nestin-positive growth cones by this technique, the very low number of definable growth cones with traceable axons per electroporated brain made quantification from this preparation not feasible. Therefore, we looked at all the growth cones we could identify to see that they were qualitatively comparable to what we had observed *in vitro*.

### Nestin does not increase global cdk5 activity towards DCX

What is the mechanism for nestin-mediated enhancement of cdk5 phosphorylation of DCX? As was reported before in other systems, we observed that p35 levels were increased by nestin co-expression (Fig. 2), likely due to stabilization of p35 by nestin binding (Pallari et al., 2011; Sahlgren et al., 2003, 2006; Yang et al., 2011). This stabilization of p35 has been proposed to constitute the mechanism by which nestin promoted cdk5 activity, since more activator should result in more cdk5 activity. To test this model directly, we expressed increasing amounts of p35 in HEK293 cells by increasing the amount of p35 plasmid used in the transfection from 1 to 4 µg. We saw an increased amount of p35 expressed with more p35 plasmid used (Fig. 6A). We could thus directly compare the levels of pS297-DCX at similar p35 levels with or without nestin. Even with similar levels of p35, levels of pS297-DCX were not increased in the absence of nestin (Fig. 6A,A’, grey bars). This demonstrated that the increased p35 levels were not sufficient to account for the increased DCX phosphorylation by cdk5 observed with nestin co-expression. Nestin thus did not increase global cdk5 activity, but selectively increased cdk5 activity towards DCX.

**Figure 6:**
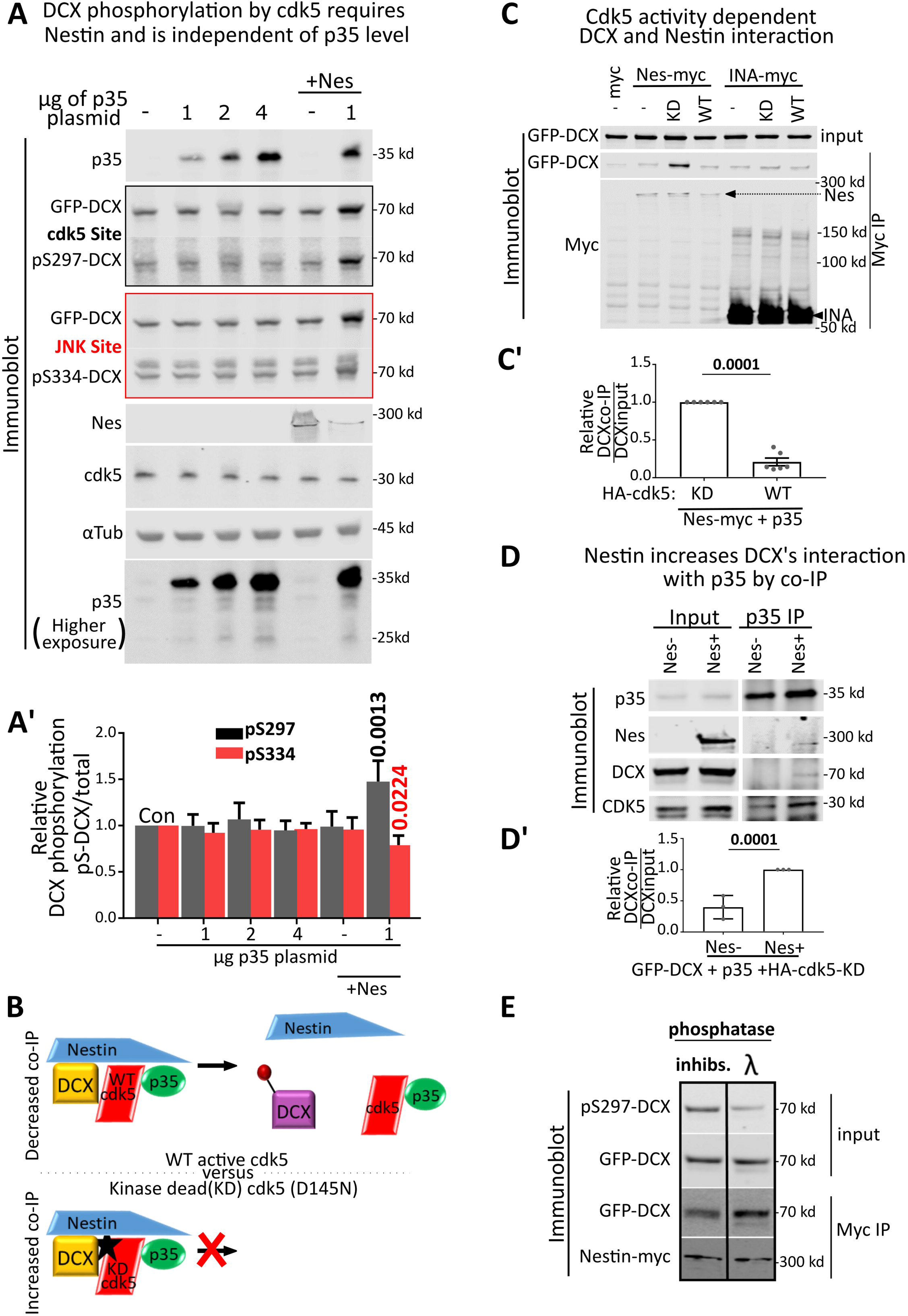
Nestin scaffolds cdk5 and DCX to promote DCX phosphorylation. A. Variable amounts of p35 plasmid (0, 1, 2, 4 µg) were transfected to increase levels of p35 expression in the absence of nestin. In addition, nestin was co-expressed with 0 or 1 µg p35 plasmid (+Nes lanes). GFP-DCX was expressed in every condition. Levels of pS297-DCX and pS334-DCX were determined by Western blotting. Levels of GFP-DCX, p35, cdk5 are also assayed. A longer exposure of the p35 blot is shown to illustrate negligible levels of the proteolytic product p25 form of p35. A’. Densitometry analysis of A. Grey bars are ratio of pS297-DCX/total GFP-DCX, red bars are ratio of pS334-DCX/total GFP-DCX. Error bars are SD. Statistical analysis was one way ANOVA with Dunnet’s correction. Significant pairwise p-values compared to control (Con) are indicated in black for pS297 and in red for pS334. All other pairwise comparisons to control were not significant and are not shown. N=4 independent experiments. *p* values of less than 0.05 are indicated in bold. B. Experimental model for using the kinase dead (KD) cdk5-D145N to trap substrate: If a quaternary complex of cdk5/p35/DCX/nestin exists, the kinase-dead cdk5 is predicted to trap all components of the complex which can now be co-IPed. WT cdk5, on the other hand, more transiently associates with its substrate DCX and does not form a stable complex (as is typical for kinases and their substrates). C. KD cdk5 increases efficiency of nestin pulldowns for DCX. GFP-DCX and p35 was co-transfected with myc EV, nes-myc, or INA-myc together with HA-cdk5-WT or HA-cdk5-KD, as indicated above the lanes, and IPed with anti-myc antibody. GFP-DCX inputs (top blot) were compared to amounts of immunoprecipitated GFP-DCX after anti-myc IPs (bottom two blots). C’. The amount of DCX after nestin IP is shown as a ratio of the co-IP to input DCX. HA-cdk5-KD was compared to HA-cdk5-WT after nestin IP. This ratio was set to 1 for cdk5-KD. With cdk5-KD, DCX is trapped in a stable complex with nestin. Error bars are SD, and statistics were unpaired t-test. N= 6 independent experiments. *p* values of less than 0.05 are indicated in bold. P35 co-IPs more DCX when nestin is co-expressed. Transfection conditions are indicated above each lane. HA-cdk5-KD is expressed in each condition to stabilize the kinase/substrate/scaffold complex. Input levels in the lysate (left panel) are compared to levels of GFP-DCX IPed with anti-p35 antibody (right panel). Nestin increases the amount of DCX that co-IP’s with p35. D’. Relative densitometry analysis of DCX in p35 co-IP compared to DCX in input. Error bars are SD, and statistical analysis was t-test with Welch’s correction. N= 3 independent experiments. Lambda (λ) phosphatase effectively reduces pS297 levels of GFP-DCX as detected in input lysates. Nestin-myc can co-IP more GFP-DCX when dephosphorylated with lambda phosphatase, compared to untreated samples containing phosphatase inhibitors (inhibs.). Binding was performed *in vitro* using overexpression lysates. A representative blot (of 4 independent experiments) is shown.

### Nestin scaffolds cdk5/p35 for DCX phosphorylation

We next asked how nestin can increase phosphorylation of DCX selectively. We hypothesized that nestin acted to scaffold the active kinase (cdk5/p35) with the substrate by binding to both proteins simultaneously (see diagram top panel Fig. 6B). Typically, interactions between a kinase and its substrate are transient, and phosphorylation of the substrate results in dissociation of the kinase from the substrate (Goldsmith et al., 2007; Manning & Cantley, 2002). Mutated kinases which are “kinase-dead” (KD) form more stable complexes with their substrates, thus “trapping” the substrate bound to the kinase. This will result in an increase in co-IP efficiency of the substrate with the kinase (bottom panel Fig. 6B). In order to test for a multimeric complex of nestin with cdk5/p35 and DCX, we compared the extent of DCX-nestin complex formation in cells co-expressing either HA-cdk5-WT or HA-cdk5-D145N (= kinase-dead KD) with p35 and nestin (Fig. 6C). We could co-IP significantly more DCX with nestin-myc (using anti-myc IP) when the substrate-trapping cdk5-KD was co-expressed, compared to low co-IP efficiency with cdk5-WT (Fig. 6C’). As a control, co-expression of INA-myc, which does not interact with p35 or DCX, resulted in only background levels of GFP-DCX co-IP in all conditions (Fig. 6C). These results suggested that nestin was acting as a scaffold for cdk5 and DCX.

To more directly test this idea, we pulled down the substrate-trapped cdk5-KD complex via p35 IP, to test for differences in DCX co-IP in the absence or presence of nestin (Fig. 6D). We found that nestin co-expression significantly increased the amount of DCX recovered in the co-IP by anti-p35 antibody, suggesting a complex of p35-cdk5-KD with DCX and nestin (Fig. 6D, 6D’). Nestin did not affect the abundance of cdk5/p35 heterodimer, as there was not a significant difference in the amount of cdk5 recovered by anti-p35 IP in the absence or presence of nestin (normalized average of 1 (no nestin) compared to 1.125 (with nestin), p=0.88, n=3).

Our model predicts that nestin would bind with higher affinity to de-phosphorylated DCX (see diagram top panel Fig. 6B). To test this, we treated GFP-DCX containing lysate, prepared from transfected HEK293 cells, with Lambda (λ) phosphatase, or with phosphatase inhibitors (inhibs.), to increase dephosphorylated or phosphorylated DCX, respectively (Fig. 6E, inputs). These crude lysates were mixed with nestin-myc containing lysates, and nestin-myc was IP’ed. We found that the sample that had been treated with the exogenous λ phosphatase (containing more de-phosphorylated DCX) was co-IP’ed more efficiently by nestin-myc then the sample that contained a higher level of pS297-DCX (Fig. 6E).

Thus, we propose that nestin acts as a selective scaffold, which can bind to cdk5/p35 and DCX simultaneously, and this results in increased efficiency of WT cdk5-mediated phosphorylation of DCX (Fig. 11). Dephosphorylation of DCX was shown by others (Bielas et al., 2007; Shmueli et al., 2006) to be dependent on the recruitment of PP1 to DCX by the actin binding protein SPN. We confirmed this observation (Fig. 2A,A’). Thus, we propose that the level of DCX phosphorylation is determined by the balance between these opposing reactions (see top half of model in Fig. 11), and that phosphorylation would be more dominant when nestin is present. We thus envision nestin to act like a “gain control” in growth cones, which sensitizes DCX to cdk5 phosphorylation.

### The T316D mutation in nestin affects p35 and DCX binding as well as DCX phosphorylation, but not incorporation into vimentin filaments

We previously showed that nestin expression is associated with small compact GCs, whereas nestin negative-neurons have larger GCs (Bott et al., 2019). Is the effect of nestin on GC morphology dependent on its ability to scaffold cdk5/p35/DCX? To test this, we aimed to create nestin mutants that lack cdk5/p35 binding. Previous work showed that the threonine at position 316 (T316) on nestin is an important regulatory site for cdk5/p35 binding and for cdk5-dependent effects of nestin (Su et al., 2013; Yang et al., 2011). The T316 site can be phosphorylated, which affects binding to p35. The T316A phospho-dead mutant had increased binding to p35 in *in vitro* binding assays of purified recombinant protein (Yang et al., 2011). In addition to making the T316A phospho-dead nestin mutant, we created a phospho-mimetic mutant (T316D-Nes), reasoning that it might be unable to bind p35. To test p35 binding of the T316 mutants, we used a co-IP strategy similar to that used in Figure 1 (Fig. 7A). In our hands, the T316A-Nes mutant did not show a consistent increase in p35 binding. T316D-Nes, on the other hand, was greatly deficient in p35 binding (Fig. 7A,A’). We then tested if these nestin mutant proteins still formed a complex with DCX. We performed IPs with WT nestin or nestin mutants when GFP-DCX was co-expressed. Surprisingly, T316D-Nes exhibited decreased binding to DCX (Fig. 7A, A’), whereas T316A-Nes still co-IP’ed DCX. We note that T316D-Nes levels appeared consistently lower in the lysates compared to WT nestin or T316A-Nes.

**Figure 7:**
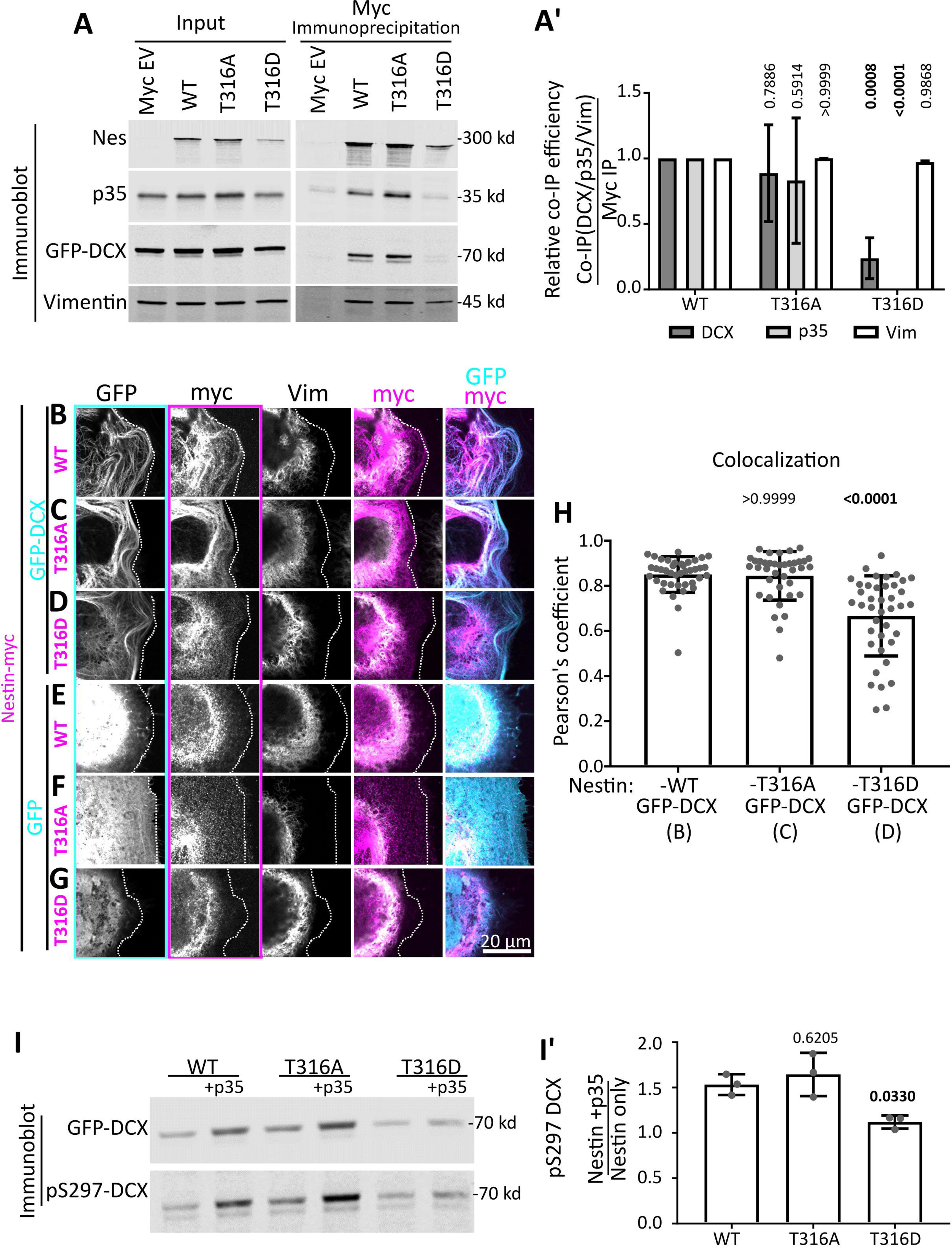
The T316D mutation in nestin affects p35/DCX binding and DCX phosphorylation, but not incorporation into vimentin filaments. A. Point mutants in nestin (T316A - phospho dead or T316D - putative phospho mimetic) were tested for binding to p35, DCX, and vimentin by co-IP from HEK293 cells. Inputs (left blot) and corresponding co-immunoprecipitation of Vimentin, GFP-DCX, and p35 after anti-myc IP (right blot) are shown. [Note, the p35 and DCX/vimentin panels are combined from the same experiment, carried out from two different plates of cells. The blots are qualitatively similar for both sets, and were combined into a single figure for simpler viewing.] A’ Densitometry quantification of co-IP efficiency (co-IP / IP) from the blots in A. T316D-nestin pulls down greatly reduced levels of p35 and DCX compared to WT nestin. Co-IP of p35 and DCX by T316A-nestin is not significantly different compared to WT nestin. Each mutant co-IPs vimentin similarly to WT. Means of n=3 experiments for each IP condition is shown, and error bars are SD. Data was analyzed with parametric two-way ANOVA test with Tukey’s correction, and p-values shown above each condition compared to the corresponding WT-Nestin co-IP. P-values less than 0.05 are in bold and are considered significant. B-G. Confocal images of COS-7 cells transfected with GFP-DCX (C-E) or GFP (F-H) co-transfected with WT nes-myc (C,F) or mutant nestin (T316A: D,G. T316D: E,H), as indicated. The transfected cell is outlined with a dotted line. Individual black and white channels are shown, as well as merges with GFP/myc and myc/vimentin channels. Outlines of black and white images correspond to the color used in the composite - images. WT nestin and the 316A-nestin mutant co-localize with DCX MT bundles. However, nestin is associated with vimentin when expressed without DCX or when the DCX-binding deficient T316D-nestin is expressed, even in the presence of DCX. H. Pearson’s coefficient of conditions represented in B, C, and D (as indicated below the x-axis labels. N=38-40 cells from 3 independent experiments are shown with the mean, and error bars are SD. Note, the WT nestin with DCX condition has the same values from figure 4F. Data was analyzed with non-parametric Kruskal-Wallis one way ANOVA test with Dunn’s correction, with each p-value shown above each condition compared to GFP-DCX with WT Nes-myc (1^st^ lane). P-values less than 0.05 and are in bold and are considered significant.. The Pearson coefficient mean for Nes-T316A with DCX is not significantly different that Nes-WT with DCX. However, Nes-T316D with DCX has a significantly (p=0.0001) lower Pearson coefficient mean than Nes-WT with DCX. I. Nestin-dependent cdk5-phosphorylation of DCX in HEK293 cells. I’. Densitometry quantification of the blot in B showing the relative levels of pS297-DCX phosphorylation with expression of each mutant. Each value represents the relative phosphorylation of DCX when expressed with nestin and p35, compared to just nestin expression. T316D-nestin does not augment p35/CDK5-mediated DCX phosphorylation. Error bars are SD, and each condition was compared to WT nestin condition (one way ANOVA with Dunnett’s correction). N= 3 independent experiments. Bold *p* values are less than 0.05 and are considered significant.

In order to test the ability of T316 nestin mutants to associate with DCX by an additional assay, we also used the COS-7 cell immunofluorescence assay to determine if nestin mutants were still recruited to MTs by DCX. In agreement with the co-IP studies, both WT nestin and T316A-Nes did not have significantly different colocalization means with GFP-DCX MT bundles, when quantified with Pearson’s coefficient (Fig. 7H). T316D-Nes, on the other hand, localized more similarly to vimentin and did not appear enriched with DCX-decorated MTs (Fig. 7B-D). T316D-Nes had significantly (p=0.0001) lower Pearson’s coefficient mean when compared to WT nestin (Fig. 7H).

Our scaffolding hypothesis proposes physical associations between DCX, p35/cdk5, and nestin. We thus reasoned that a nestin mutant deficient in p35 and DCX binding should also be deficient in promoting cdk5-mediated phosphorylation of DCX. We tested if the pS297-DCX levels were altered in cells expressing WT or T316 nestin mutants (Fig. 7I). T316A-Nes + p35 resulted in increased phosphorylation of DCX similar to WT nestin + p35, whereas T316D-Nes + p35 did not result in increased phosphorylation of DCX (Fig. 7I, quantified in 7I’). This result is consistent with our scaffolding hypothesis.

Since nestin forms heteropolymers with vimentin, we also tested if the nestin T316 mutants still interacted with vimentin. The T316 site is just outside of the IF polymerization rod domain, so modification of this site is not expected to interfere with IF polymerization. Indeed, we observed WT nestin, T316A-Nes, and T316D-Nes co-IP’ing vimentin similarly (Fig. 7A, A’), arguing that they still formed homo-oligomers/polymers with vimentin. In agreement, these mutants appeared to co-distribute with endogenous vimentin filaments similarly to WT nestin when co-expressed in COS-7 cells (Fig. 7E-G). We thus conclude that T316D-Nes binds less well to both p35 and DCX whereas T316A-Nes still binds p35 and DCX, and that neither mutation grossly disrupts vimentin binding. We use the T316D-Nes mutant to test the role of the nestin-p35-DCX complex in neurons below.

### Binding to p35 and/or DCX is required for nestin’s effect on growth cone size

We previously reported that reducing neuronal nestin levels leads to larger GCs (Bott et al., 2019). We thus wondered if overexpression of WT nestin would do the opposite, i.e. lead to smaller GCs. We expressed myc-nestin in cultured cortical neurons for 2 days along with GFP to allow for detection of transfected cells. At this stage, the majority of neurons are nestin-negative (Bott et al., 2019), allowing us to assess the effects of re-expressing nestin. Overexpression of WT nestin-myc resulted in statistically smaller GCs on axons (arrows) with fewer filopodia, which is the opposite phenotype of downregulating nestin. Overexpression of INA-myc did not change GC size or filopodia number compared to myc-EV controls (Fig. 8A, B, B’). Next we asked if p35/DCX binding by nestin was required for the effect on GC size. T316A-Nes, which retained binding to p35 and DCX (Fig. 7A) and resulted in WT levels of DCX phosphorylation (Fig. 7B), resulted in small GCs with fewer filopodia, similar to WT nestin (Fig. 8A, B, B’). However, the T316D-Nes mutant, which did not interact with p35 or DCX (Fig. 7A) and did not result in increased DCX phosphorylation (Fig. 7B), did not significantly alter GC size or filopodia number (Fig. 8A, B, B’). Therefore, this result suggests that binding to p35 and/or DCX is required for nestin’s effect on GC size.

**Figure 8:**
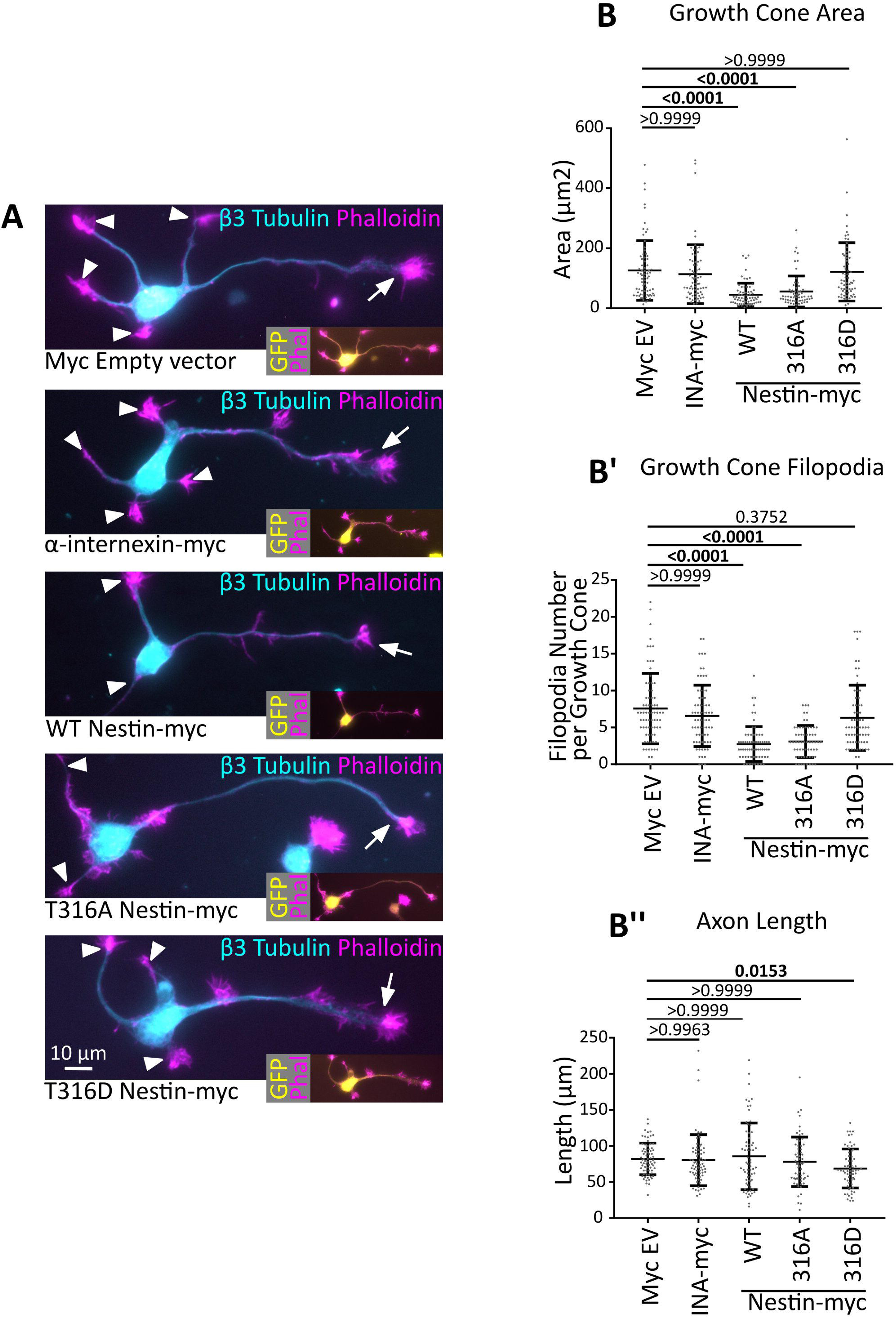
Binding to p35 and/or DCX is required for nestin’s effect on growth cone size. A. Representative images of DIV2 cortical neurons expressing nestin or T316-nestin mutants compared to empty vector control or INA-myc. β3 tubulin and phalloidin were used to visualize MTs and F-actin, respectively. Axonal GCs are indicated with arrows, and secondary neurites (dendrites), are labeled with arrowheads. Brightness was adjusted for each image for easier viewing. Insets show transfected neurons, identified by GFP expression. B, B’, and B’’. Quantification of neuronal morphology in neurons expressing different nestin constructs (as shown in A). N= 72 to 61 cells counted per condition in 3 different independent experiments. Data was analyzed with non-parametric Kruskal-Wallis one way ANOVA test with Dunn’s correction. Pairwise p-values are shown for comparisons to myc EV (control). WT nestin and T316A-nestin decrease growth cone area and filopodia number, but T316D-nestin does not. *p* values of less than 0.05 are indicated in bold.

### Nestin does not have any effect on GC size in the absence of its downstream effector DCX

Since nestin selectively bound to DCX and affected its phosphorylation, we hypothesized that the effects of nestin overexpression on GCs were dependent on DCX. Therefore, we tested if nestin overexpression would affect GCs in the absence of DCX. We thus overexpressed nestin-myc, myc-EV, or myc-INA in WT or DCX KO neurons and determined GC size and filopodia number. We again observed decreased axonal GC size (arrows) and decreased filopodia number specifically with nestin-myc expression (Fig. 9A,B,B’). However, in DCX null neurons, no decrease in GC size or filopodia number was seen with nestin-myc expression (Fig. 9A,B,B’). When GFP-DCX was reintroduced together with nestin-myc into DCX null neurons, the nestin effect on GC size and GC filopodia was rescued (Fig. 9A,B,B’). Axon length was not affected (Fig. 9B’’). We thus conclude that nestin acts via DCX to influence GC size.

**Figure 9:**
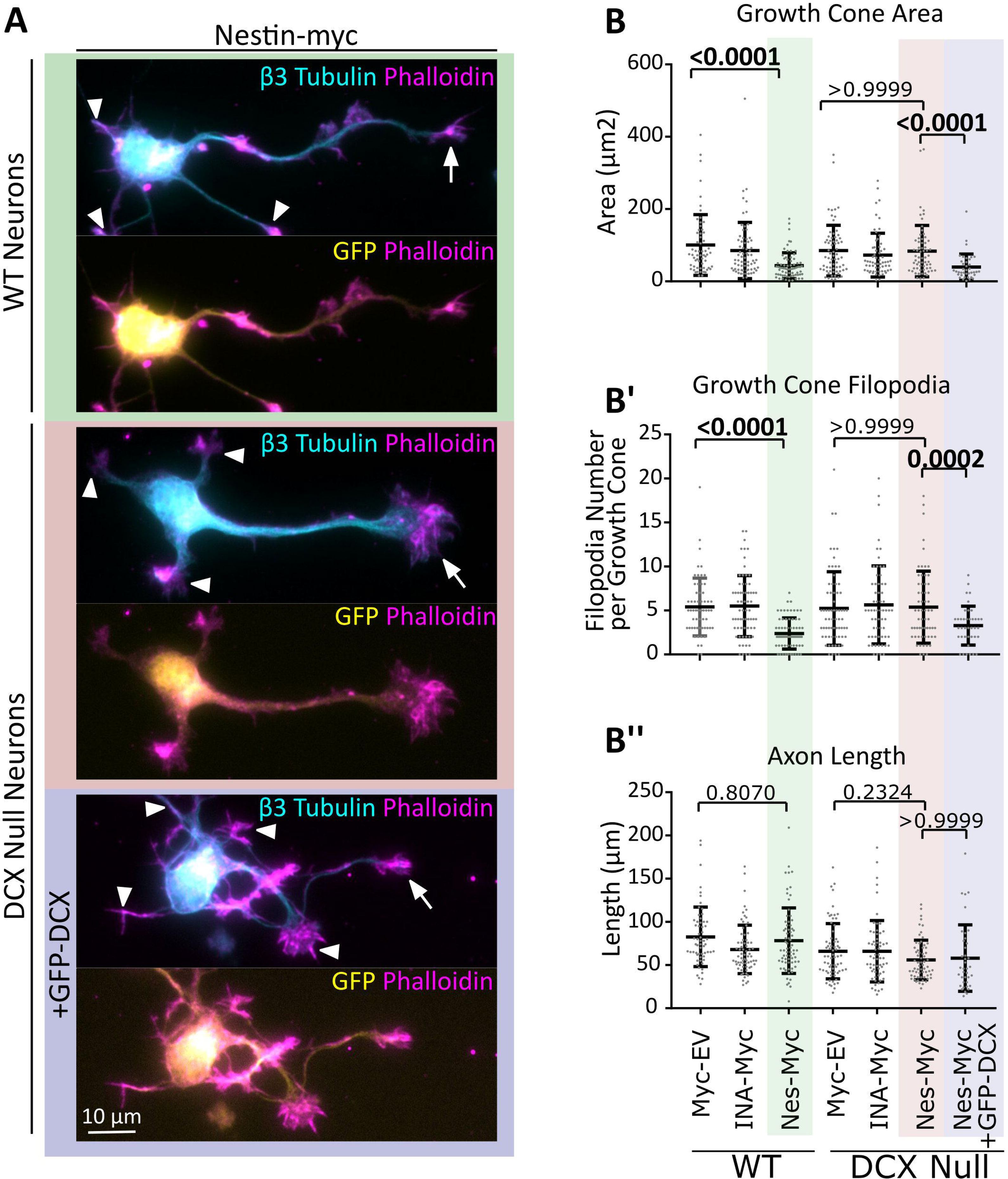
In the absence of DCX, nestin overexpression does not result in small growth cones. A. Representative images of WT or DCX null E16 cortical neurons (as labeled) expressing nes-myc with GFP or nes-myc with GFP-DCX after 48 hours in culture. WT neurons transfected with nes-myc show smaller axonal GCs (arrows) whereas DCX KO neurons transfected with nestin-myc do not show smaller GCs. When GFP-DCX is co-transfected into DCX KO neurons, GCs are smaller. Axon growth cones are indicated with arrows. The growth cones of secondary neurites (dendrites) where not considered and are indicated with arrowheads. B’,B”, B”’. Quantification of morphological characteristics. The background shading color behind the images in A correspond to the conditions shaded in the graphs B’, B’’, and B’’’ for easier interpretation. B’, B’’, and B’’’ shows the quantification of morphological measurements from transfected neurons: growth cone area (B’), filopodia number (B”), and axon length (B”’). Error bars are SD. Kruskal-Wallis one way ANOVA non-parametric test was used. P-values are only shown for selected statistical comparisons to keep the graph less cluttered. N= 40 to 70 cells counted in 3 different independent experiments.

### DCX is a cdk5 substrate downstream of Sema3a, and nestin-sensitivity to Sema3a is absent in DCX null neurons

We previously showed that 1DIV cortical WT neurons respond to Sema3a by retracting their axon growth cone filopodia and that this response is blunted when nestin is downregulated. We wondered if DCX was also downstream of nestin for Sema3a responsiveness. S297-DCX phosphorylation has been reported to be induced by Sema3a in DRG neurons (Ng et al., 2013), but this has not been shown in other neuronal cell types. We treated DIV1 E16 cortical mouse neurons with vehicle or 1 nM Sema3A for 5, 15, or 30 minutes and determined the levels of DCX phosphorylation.

We could detect phosphorylation of the S297-DCX epitope in cortical neurons, which was significantly increased after Sema3a treatment (Fig. 10A,A’ grey bars). The JNK site pS334 was not increased after Sema3a treatment (Fig. 10A,A’ red bars). pS297DCX, but not pS334DCX levels, were reduced by roscovitine, consistent with activation of cdk5 downstream of Sema3a (Fig. 10A,A”).

**Figure 10:**
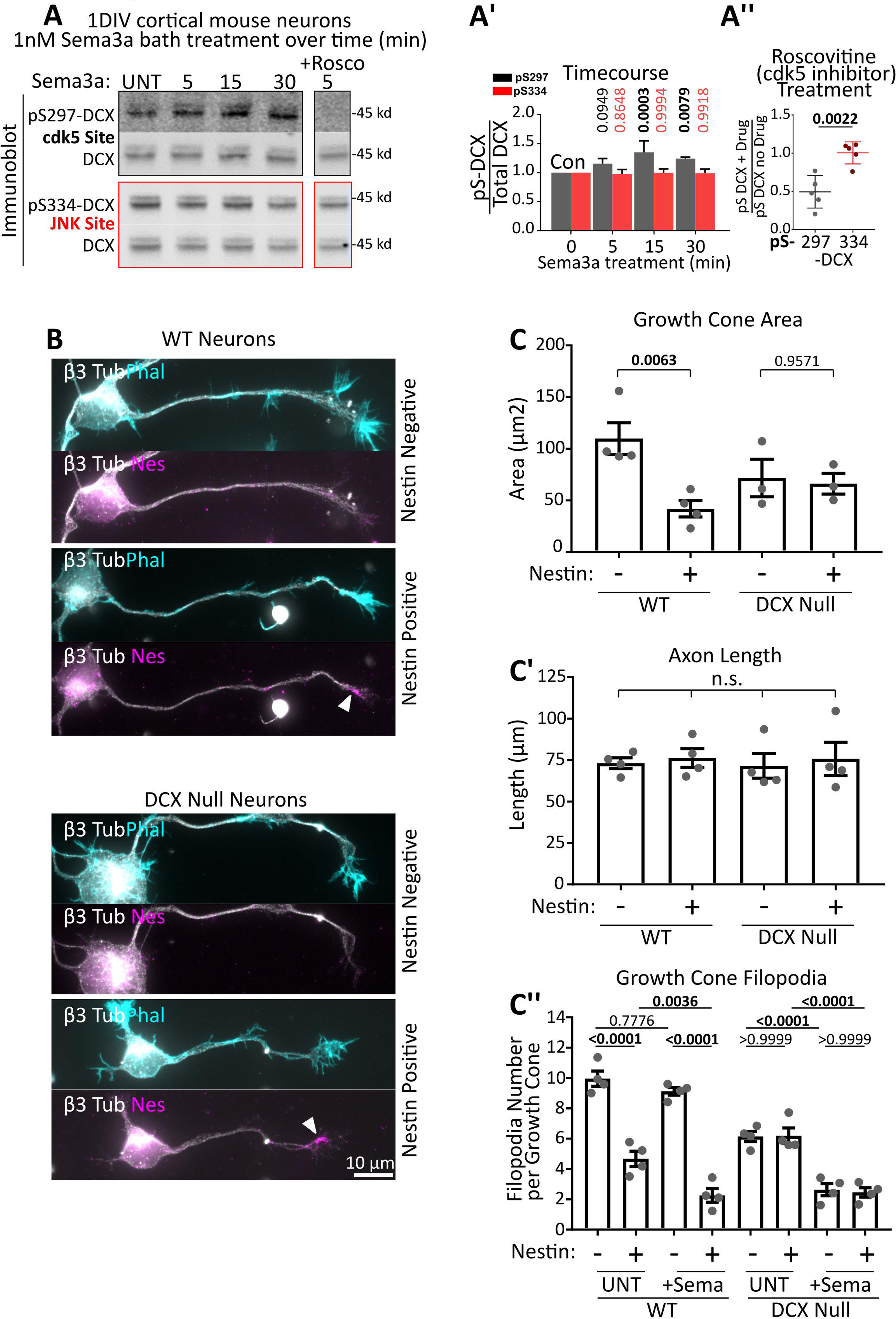
DCX is a cdk5 substrate downstream of Sema3a, and DCX null neurons respond in a nestin independent manner. A. Sema3a treatment of 1DIV cultured mouse cortical E16 neurons significantly increases DCX phosphorylation at S297, but not at S334. Rightmost lane: Neurons were treated with 1nM Sema3a for 5 min with pre-incubation of the cells with 10µM Roscovitine for 30 minutes. The inhibitor lane was from the same blot as the other lanes, but separated by intervening lanes which were removed. A’. Quantification of Sema3a time course. Blots in A were subjected to densitometry analysis, and the ratio of phospho-DCX signal relative to the total DCX is plotted. DCX phosphorylation at pS297, but not at pS334, begins quickly after Sema3a stimulation, and becomes significant at 15 and 30 minutes. Error bars are SD and statistical analysis was performed at each timepoint compared to untreated (UNT) with an ordinary one way ANOVA with Dunnett’s correction. n=5 independent experiments. *p* values of less than 0.05 are indicated in bold. A’’. Quantification of Roscovitine sensitivity (rightmost lane in A). Each value was obtained from densitometry analysis of western blots, where the relative level of p-DCX was compared to total DCX. The values in the plot are a ratio of the relative phosphorylation after inhibitor (drug) treatment compared to no inhibitor treatment. Error bars are SD and statistical analysis was performed with a two-tailed unpaired t-test. N=5 independent experiments. B. Representative images of nestin positive or negative individual neurons from WT or DCX Null primary E16 cortical neuron cultures 1DIV (stage3). Quantified in C and C’. The upper image for each condition shows phalloidin staining, and the lower image show nestin staining of the same neuron. Nestin positive growth cones are indicated with arrowheads. Both the upper and lower images have β3 tubulin shown to both visualize the cells overall morphology and to confirm neuronal identity. C, C’, C’’. Quantification of morphological characteristics of nestin-negative and nestin-positive neurons derived from WT or DCX null E16 cortex at 1DIV (stage 3). WT cells show nestin dependent decrease in growth cone size, while DCX null neurons do not. Axon length was not significantly different in any condition. When treated with 1nM Sema3a for 5 minutes, filopodial retraction in WT neurons was dependent on nestin expression. In DCX null neurons, there was a filopodia decrease after Sema3a treatment in nestin positive and negative neurons (independent of nestin). C. Statistical analysis was one way ANOVA with Sidak’s multiple comparison correction with pairs tested shown. C’ Statistical analysis was one way ANOVA with Tukey’s multiple comparison correction. C’’. Statistical analysis was one way ANOVA with Sidak’s multiple comparison correction with pairs tested shown. C, C’, and C’’ n=4-5 independent experiments with 20-30 cells counted in each set. *p* values of less than 0.05 are indicated in bold. Error bars are SEM.

**Figure 11:**
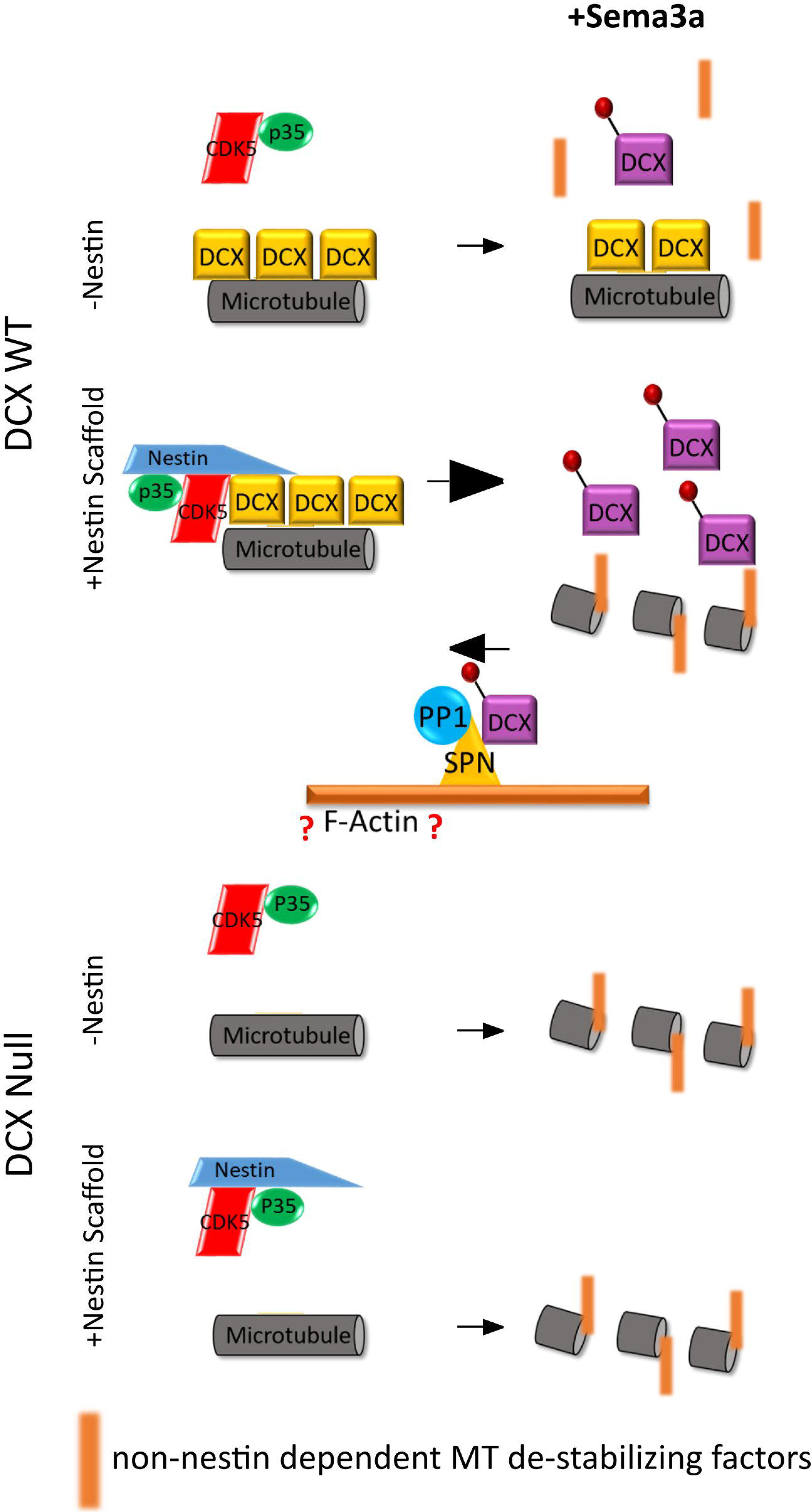
Model of nestin mediated scaffolding of Cdk5 for modulation of DCX phosphorylation. Model for how nestin increases cdk5-phosphorylation of DCX by scaffolding: in the absence of nestin (-Nestin; top) non-phospho-DCX is bound to microtubules and inefficiently phosphorylated by cdk5/p35. In the presence of nestin (+ Nestin scaffold; bottom), DCX becomes associated with cdk5/p35, increasing the efficiency of DCX phosphorylation. DCX phosphorylation results in dissociation of DCX from MTs, this allowing remodeling by additional factors (red bars). In this model, we also depict data from others that showed that the actin-associated SPN which scaffolds Protein Phosphatase 1 (PP1) potently dephosphorylates DCX. In our model, the level of phospho-DCX is determined by the relative opposing activities of the two regulatory complexes nestin/cdk5/p35 or SPN/PP1. However, in DCX null neurons (bottom half of the panel), nestin no longer has an effect on sema3a sensitivity. Without DCX MT binding and stabilization, growth cones retract and are sensitive to Sema3a independent of nestin. Nestin-mediated enhancement of Sema3a sensitivity is thus lost in DCX null neurons.

In order to determine if nestin still affected the extent of Sema3a sensitivity in the absence of DCX, we compared the Sema3a response of nestin-positive and nestin-negative neurons in WT or DCX null cortical cultures. At 1DIV a little more than half of the stage 3 neurons are nestin-positive, while the remainder are nestin-negative (Bott et al., 2019), allowing us to compare Sema3a responsiveness in the same culture when stained against nestin. In WT neurons, we again saw a nestin-dependent decrease in growth cone size in untreated cultures, whereas the growth cone size of nestin-positive and nestin-negative neurons was not significantly different in DCX null cultures (Fig. 10B, quantified in 10C), in agreement with the data in Figure 9. As we reported previously (Bott et al., 2019), we again observed a decrease in filopodia number in nestin-expressing neurons when treated with Sema3a whereas nestin-negative neurons were less Sema3a-responsive (Fig. 10C’’). When we treated DCX null neurons with Sema3a, however, we no longer observed decreased GC filopodia number in nestin-positive neurons. Nestin-positive and nestin-negative neurons derived from DCX null mice showed the same Sema3a sensitivity (Fig. 10C’’), indicating that DCX is required for nestin-dependent regulation of GC size and Sema3a sensitivity. Axon length was not significantly different (Fig. 10C’).

In our model (Fig. 11), we propose that basally, non-phospho DCX binding to MTs stabilizes the MTs, and in the absence of nestin-scaffolded cdk5 phosphorylation, DCX provides resistance or buffering against Sema3a-mediated growth cone collapse and filopodial retraction. Activation of the pathway in nestin-positive WT neurons allows for DCX phosphorylation and removal of the MT stabilizing effects of DCX. When DCX is absent and is not there to provide resistance to Sema3a, growth cones are responsive to Sema3a in a non-nestin dependent manner.

## Discussion

A number of IF subtypes are expressed in the brain, but their cell biological functions during development are incompletely understood (Yuan & Rao, 2012; Bott & Winckler, 2020). One of the earliest expressed IFs in the developing brain is nestin, but little is known about nestin’s function, even in NPCs where it is highly expressed (Mohseni et al., 2011; Park et al., 2010; Wilhelmsson et al., 2019). We previously found that nestin protein expression persists at low levels in early neurons and that nestin expression influences growth cone size and responsiveness to Sema3a in a cdk5-dependent manner (Bott et al., 2019), but the molecular pathways for this phenotype were not known. Nestin is known to directly interact with the cdk5/p35 heterodimer. We thus hypothesized that nestin might also affect the activity of cdk5 in neurons, similarly to what occurs during neuromuscular junction development (Yang et al., 2011). Hence, we screened known neuronal cdk5 substrate cytoskeletal proteins for nestin binding and identified DCX as a novel nestin-associated protein that is preferentially phosphorylated in the presence of nestin and cdk5/p35. Mechanistically, we find that nestin selectively promotes robust phosphorylation of DCX by cdk5/p35 through scaffolding. We show that DCX and nestin co-localize in the distal axon and growth cone, and that DCX is required for the effects of nestin on GC morphology and Sema3a sensitivity. In summary, our results uncover a novel signaling role for neuronal nestin in regulating GC morphology specifically via scaffolding the cdk5 substrate DCX.

### DCX is a substrate for nestin-scaffolded cdk5 in neurons

Earlier work identified nestin as a p35 binding protein and showed that nestin modulated both the level and localization of cdk5 activity in C2C12 myotubes (Yang et al., 2011). *In vivo*, nestin depletion phenocopies cdk5 deletion or roscovitine treatment in maintaining acetylcholine receptor clustering in agrin (AGD) mutants (Lin et al., 2005; Mohseni et al., 2011; Yang et al., 2011). However, the effect of nestin depletion on cdk5 activity (promoting or inhibiting) has varied between reports and assay systems and has been a point of contention (Huang et al., 2017; Lindqvist et al., 2017; Pallari et al., 2011; Sahlgren et al., 2006; J. Wang et al., 2017; Yang et al., 2011; Zhang et al., 2018). Since the relevant endogenous cdk5 substrate in myoblasts remains unknown, exogenous substrates (such as recombinant H1 histone protein) are commonly used for phosphorylation assays. The different interpretations are thus likely due to the use of exogenous substrates in the phosphorylation assays. Our work for the first time identifies a biologically relevant substrate for nestin-scaffolded cdk5, namely DCX (Fig. 1 and 3). If DCX is also the relevant substrate in myoblasts is not known, but it is intriguing that DCX null mice also have developmental defects at the neuromuscular junction, as do human lissencephaly patients with DCX mutations (Bourgeois et al., 2015). Our work suggests that nestin does not modulate cdk5/p35 kinase activity per se since nestin did not affect several other cdk5 substrates, nor was cdk5/p35 heterodimer formation affected (Fig.1 and 6D). Rather, we propose that in neurons nestin selectively increases cdk5-mediated DCX phosphorylation by providing a scaffold. We note that DCX may not be the only cdk5 substrate affected by nestin, especially in other tissues, but we focused our study on known neuronal cdk5 substrates to which specific antibodies to phospho-epitopes were available. Analysis of changes in the phosphoproteome of nestin null neurons will be required to more completely assess all the nestin-dependent phospho-proteins

### DCX as a downstream effector of nestin: a model for regulation of GC size

How does nestin affect GC size via DCX? Three major processes are necessary for axon growth and guidance - protrusion, engorgement, and consolidation (Dent & Gertler, 2003). Protrusions towards positive guidance cues are mediated by actin-based filopodia and lamella which serve as templates for subsequent MT infiltration and growth. These MTs stabilize the new actin-based structures and allow for transport of cell components through vesicular trafficking which leads to the filling and engorgement of the advancing growth cone. The absence of DCX results in decreased stability and extent of protruding MTs (Jean et al., 2012). Lastly, the proximal part of the growth cone is consolidated, and MTs are bundled into new axon shaft. MT-binding proteins play essential roles in regulating the assembly dynamics and bundling extent of MTs in a spatially regulated manner and are critical regulators of axon growth. Phosphorylation and dephosphorylation are known to spatially regulate the activities of many MAPs, including DCX. We propose that nestin affects GC size by promoting the phosphorylation of DCX in the distal axon. Dephosphorylation of DCX has already been implicated in the consolidation step of axon growth (Bielas et al., 2007). Specifically, Bielas et al. (2007) showed that cdk5-mediated phosphorylation of DCX is reversed by the phosphatase PP1 which is scaffolded by the DCX-binding protein SPN (Tsukada et al., 2006). In extension of these earlier models, we propose that nestin scaffolds cdk5 and thereby promotes the phosphorylation of DCX - thus completing the regulatory cycle of DCX phosphorylation and dephosphorylation (Fig. 11). The regulation of the phosphorylation of DCX is thus driven by the balance of nestin/cdk5/p35 to increase phosphorylation and SPN/PP1 to decrease it - perturbation of either process results in abnormal MT organization (Bielas et al., 2007; Dehmelt & Halpain, 2007; Schaar et al., 2004). In our model, the level of phospho-DCX would be determined by the relative opposing activity of the two differently localized regulatory complexes (Fig. 11).

On a molecular level, we predict that the presence of nestin in growth cones would result in more pS297-DCX, which has a decreased affinity for MTs. We were unable to stain neurons with the pS297-DCX antibody reliably and could thus not test this idea directly. Since DCX binding to MTs stabilizes them (Moores et al., 2004), growth cones with more pS297-DCX may have intrinsically less stable or more dynamic MTs. This notion is consistent with previous findings that GC size reflects MT dynamics. For instance, smaller growth cones have intrinsically more dynamic MTs than larger growth cones (Kiss et al., 2018). In addition, treatment with taxol, which stabilizes MTs, results in larger GCs (Gumy et al., 2013). Thus we propose that a smaller growth cone represents a GC with more robust kinase/phosphatase activity which would result in more rapid cytoskeletal dynamics, morphological turnover, and GC sculpting via changes in the phosphorylation of cytoskeletal associated proteins, including DCX. A smaller GC would thus correlate with a more robust response to a given relevant guidance cue as it is able to remodel the existing cytoskeleton more rapidly.

### A novel role for intermediate filaments in axon growth

IFs have previously not been thought to be important for axon formation, much less axon guidance. Early studies had concluded that IFs were absent from the most distal and peripheral regions of the growth cone, and thus were ruled out early on as relevant players in regulating axon targeting (Dent & Gertler, 2003). However, the higher resolution and more sensitive imaging methods used here (Fig. 5B), reveal IFs in some peripheral regions of the GC, suggesting this question should be revisited. Indeed, nestin mRNA has been consistently found to be enriched in neurites relative to the cell body (Kügelgen & Chekulaeva, 2020). It remains thus an open question whether or not some IF proteins can affect axon guidance or outgrowth in some ways *in vivo*.

Nestin-containing IFs are indeed well localized to influence growth cone behavior. In support of our *in vitro* data, DCX and nestin co-express *in vivo* while neurons are in the stage of axon positioning and orientation (Namba et al., 2014)(Fig. 5 C,D) which are cdk5- and Sema3a-dependent processes (Oshima et al. 2007; Polleux et al., 1998). Our own work shows striking accumulation of nestin in the distal axon shaft and even in some peripheral regions of the axonal growth cone (Fig. 5). Nestin co-localizes with a subset of DCX, but is not identically distributed. This raises interesting future questions about why nestin is not present in dendrites, and if this distribution is related to known differences in behavior of axonal and dendritic growth cones (i.e. differential guidance cue sensitivity) (Wang et al., 2014).

### Summary

This work highlights how the IF composition of neurons can alter intracellular signaling pathways and thus alter the cell’s morphology. IFs have long been implicated as mediators and regulators of “cytoskeletal crosstalk” (Chang & Goldman, 2004; Michalczyk & Ziman, 2005), and our findings here represent an example of cytoskeletal crosstalk between IFs and MTs and their associated proteins. Our previous work has found that nestin expression correlated with small GC size and greater responsiveness to Sema3a. Based on our new work, we implicate modulation of DCX phosphorylation by cdk5 as a relevant molecular pathway downstream of neuronal nestin function. We thus posit that nestin sensitizes growth cones to cdk5 activity by scaffolding DCX. Based on our data, we propose the model that nestin affects growth cone morphology as a kinase regulator. In addition to this signaling role, nestin could also play a structural role as a cytoskeletal filament, but more work will be required to more fully elucidate nestin’s role in neurons during development.

## Acknowledgements

From the University of Virginia, we thank the Deppman lab for TUJ1 antibody, Laura Digilio for assistance with editing and Airyscan imaging expertise, Ashley Mason for assistance with editing, the Kipnis lab for use of Amaxa 4d electroporator. We also thank Lawrence Brass (University of Pennsylvania) for Myc-Spinophilin expression construct, Bruce Lahn’s lab (University of Chicago) for generously providing mouse nestin cDNA, Dr. Martin Balastik (Cech Academy of Sciences) for generously providing the CRMP2 reagents, Clay Hazlett from Nikon for assistance with the Nikon TiE enhanced confocal imaging, and Dr. Judy Liu (Brown University) for the DCX KO mice. This work was supported by NIH grant R01NS081674 (to BW). CB was supported by an institutional NIH training grant (T32 GM008136).

CB and BW conceived and coordinated the study, interpreted the data, and wrote the paper. CB devised, performed, and analyzed all experiments except for *in utero* electroporations. JK and KYK devised and performed CRISPR *in utero* electroporation experiments (Fig. 2C-F). LPM performed blinded analysis of colocalization and morphology data sets. CY provided critical expertise for experimental design, execution, and analysis.

## References

1. Bahi-Buisson, N., Souville, I., Fourniol, FJ., Toussaint, A., Moores, CA., Houdusse, A., Lemaitre, JY., Poirier, K., Khalaf-Nazzal, R., Hully, M., Leger, PL., Elie, C., Boddaert, N., Beldjord, C., Chelly, J., Francis, F. (2013). New insights into genotype-phenotype correlations for the doublecortin-related lissencephaly spectrum. Brain: A Journal of Neurology, 136(Pt 1), 223–244.

2. Benson, DL., Mandell JW., Shaw, BG. (1996). Compartmentation of alpha-internexin and neurofilament triplet proteins in cultured hippocampal neurons. Journal of Neurocytology, 25(3), 181–196.

3. Bielas, SL., Serneo, FF., Chechlacz, M., Deerinck, TJ., Perkins, GA., Allen, PB., Ellisman, MH., Gleeson, JG. (2007). Spinophilin Facilitates Dephosphorylation of Doublecortin by PP1 to Mediate Microtubule Bundling at the Axonal Wrist. Cell, 129(3), 579–591.

4. Bott, C. J., Johnson, C. G., Yap, C. C., Dwyer, N. D., Litwa, K. A., & Winckler, B. (2019). Nestin in immature embryonic neurons affects axon growth cone morphology and Semaphorin3a sensitivity. Mol Biol Of Cell, 30(10), 1214–1229.

5. Bott, C.J. and Winckler, B. (2020). Intermediate filaments in developing neurons: Beyond structure. Cytoskeleton. 2020;1–19.

6. Bourgeois, F., Messéant, J., Kordeli, E., Petit, J. M., Delers, P., Bahi-Buisson, N., … Legay, C. (2015). A critical and previously unsuspected role for doublecortin at the neuromuscular junction in mouse and human. Neuromuscular Disorders, 25(6), 461–473.

7. Brown, A. (2013). Axonal Transport. In Neuroscience in the 21st Century (pp. 255–308).

8. Chang, L., & Goldman, R. D. (2004). Intermediate filaments mediate cytoskeletal crosstalk. Nature Reviews. Molecular Cell Biology, 5(8), 601–613.

9. Cheung, Z. H., & Ip, N. Y. (2012). Cdk5: A multifaceted kinase in neurodegenerative diseases. Trends in Cell Biology, 22(3), 169–175.

10. Contreras-Vallejos, E., Utreras, E., Bórquez, D. a, Prochazkova, M., Terse, A., Jaffe, H., Toledo, A., Arruti, C., Pant, HC., Kulkarni, AB., González-Billault, C. (2014). Searching for novel Cdk5 substrates in brain by comparative phosphoproteomics of wild type and Cdk5-/-mice. PloS One, 9(3), 1–13.

11. Dehmelt, L., Halpain, S. (2007). Neurite outgrowth. A Flick of the wrist. Current Biology, 17(15), 609–611.

12. Dent, E. W., & Gertler, F. B. (2003). Cytoskeletal dynamics and transport in growth cone motility and guidance. Neuron, 40(2), 209–227.

13. Dent, E. W., Gupton, S. L., & Gertler, F. B. (2011). The growth cone cytoskeleton in axon outgrowth and guidance. Cold Spring Harbor Perspectives in Biology, 3(3), 1– 39.

14. Dhavan, R., & Tsai, L. (2001). A Decade of Cdk5. Nature, 2(October), 749–759.

15. Friocourt, G., Koulakoff, A., Chafey, P., Boucher, D., Fauchereau, F., Chelly, J., & Francis, F. (2003). Doublecortin functions at the extremities of growing neuronal processes. Cerebral Cortex, 13, 620–626.

16. Ge, W., He, F., Kim, K. J., Blanchi, B., Coskun, V., Nguyen, L., Wu, X., Zhao, J., Heng, JI., Martinowich, K., Tao, J., Wu, H., Castro, D., Sobeih, MM., Corfas, G., Gleeson, JG., Greenberg, ME., Guillemot, F., Sun, YE. (2006). Coupling of cell migration with neurogenesis by proneural bHLH factors. Proceedings of the National Academy of Sciences of the United States of America, 103(5), 1319–1324.

17. Geraldo, S., & Gordon-Weeks, P. R. (2009). Cytoskeletal dynamics in growth-cone steering. Journal of Cell Science, 122(Pt 20), 3595–3604.

18. Gleeson, J. G., Peter T, L., Flanagan, L. a., & Walsh, C. a. (1999). Doublecortin is a microtubule-associated protein and is expressed widely by migrating neurons. Neuron, 23(2), 257–271.

19. Goldsmith, E.J., R. Akella, X. Min, T. Zhou, J. H. (2007). Substrate and Docking Interactions in Ser/Thr Protein Kinases. Chemical Reviews, 107(11), 5065–5081.

20. Gumy, LF., Chew, DJ., Tortosa, E., Katrukha, EA., Kapitein, LC., Tolkovsky, AM., Hoogenraad, CC., Fawcett, JW. (2013). The Kinesin-2 Family Member KIF3C Regulates Microtubule Dynamics and Is Required for Axon Growth and Regeneration. The Journal of Neuroscience, 33(28), 11329–11345.

21. Hirasawa, M., Ohshima, T., Takahashi, S., Longenecker, G., Honjo, Y., Veeranna, Pant, H. C., Mikoshiba, K., Brady, R., Kulkarni, A. B. (2004). Perinatal abrogation of Cdk5 expression in brain results in neuronal migration defects. PNAS, 101(16), 6249–6254.

22. Huang, Y., Wu, C., Shi, G., & Wu, G. C. (2017). Nestin Serves as a Prosurvival Determinant that is Linked to the Cytoprotective Effect of Epidermal Growth Factor in Rat Vascular Smooth Muscle Cells. Journal of Biochemistry, 146(October), 307– 315.

23. Jean, D. C., Baas, P. W., & Black, M. M. (2012). A novel role for doublecortin and doublecortin-like kinase in regulating growth cone microtubules. Human Molecular Genetics, 21(26), 5511–5527.

24. Kapitein, L. C., & Hoogenraad, C. C. (2015). Building the Neuronal Microtubule Cytoskeleton. Neuron, 87(3), 492–506.

25. Kawauchi, T. (2014). Cdk5 regulates multiple cellular events in neural development, function and disease. *Development*, Growth & Differentiation, 56(5), 335–348.

26. Kiss, A., Fischer, I., Kleele, T., Misgeld, T., & Propst, F. (2018). Neuronal Growth Cone Size-Dependent and -Independent Parameters of Microtubule Polymerization. Frontiers in Cellular Neuroscience, 12(July), 1–15.

27. Kügelgen, N. Von, & Chekulaeva, M. (2020). Conservation of a core neurite transcriptome across neuronal types and species, (November 2019), 1–16.

28. Laser-Azogui, A., Kornreich, M., Malka-Gibor, E., & Beck, R. (2015). Neurofilament assembly and function during neuronal development. Current Opinion in Cell Biology, 32(2), 92–101.

29. Lin, W., Dominguez, B., Yang, J., Aryal, P., Brandon, E. P., Gage, F. H., & Lee, K. (2005). Neurotransmitter Acetylcholine Negatively Regulates Neuromuscular Synapse Formation by a Cdk5-Dependent Mechanism. Neuron, 46(5), 569–579.

30. Lin, W., Szaro, B. (1995). Neurofilaments Help maintain Normal Morphologies and Support Elongation of Neurites in Xenopus laevis Cultured Embryonic Spinal Cord Neurons. The Journal of Neuroscience, 15(December), 8331–8344.

31. Lin, W., & Szaro, B. G. (1996). Effects of Intermediate Filament Disruption on the Early Development of the Peripheral Nervous System of Xenopus laevis. Developmental Biology, 211(0251), 197–211.

32. Lindqvist, J., Torvaldson, E., Gullmets, J., Karvonen, H., Nagy, A., Taimen, P., & Eriksson, J. E. (2017). Nestin contributes to skeletal muscle homeostasis and regeneration. Journal of Cell Science, 130, 2833–2842.

33. Manning, B. D., & Cantley, L. C. (2002). Hitting the Target L the Search for Kinase Substrates. Sci STKE, 162(12), 1–5.

34. Menon, S., & Gupton, S. L. (2016). Building Blocks of Functioning Brain: Cytoskeletal Dynamics in Neuronal Development. In International Review of Cell and Molecular Biology (Vol. 3, pp. 183–245).

35. Michalczyk, K., & Ziman, M. (2005). Nestin structure and predicted function in cellular cytoskeletal organisation. Histology and Histopathology, 20, 665–671.

36. Miller, K. E., & Suter, D. M. (2018). An Integrated Cytoskeletal Model of Neurite Outgrowth. Frontiers in Cellular Neuroscience, 12(November), 1–19.

37. Mohseni, P., Sung, H-K., Murphy, A. J., Laliberte, C. L., Pallari, H.-M., Henkelman, M., Georgiou, J., Xie, G., Quaggin, SE., Thorner, PS., Eriksson, JE., Nagy, A. (2011). Nestin is not essential for development of the CNS but required for dispersion of acetylcholine receptor clusters at the area of neuromuscular junctions. The Journal of Neuroscience 31(32), 11547–11552.

38. Moores, C. A., Perderiset, M., Francis, F., Chelly, J., Houdusse, A., & Milligan, R. A. (2004). Mechanism of microtubule stabilization by doublecortin. Molecular Cell, 14(6), 833–839.

39. Moslehi, M., Ng, D. C. H., & Bogoyevitch, M. A. (2017). Dynamic microtubule association of Doublecortin X (DCX) is regulated by its C-terminus. Scientific Reports, 5245(7), 1–11.

40. Namba, T., Kibe, Y., Funahashi, Y., Nakamuta, S., Takano, T., Ueno, T., … Takeuchi, K. (2014). Pioneering Axons Regulate Neuronal Polarization in the Developing Cerebral Cortex. Neuron, 81(4), 814–829.

41. Ng, T., Ryu, J. R., Sohn, J. H., Tan, T., Song, H., Ming, G. L., & Goh, E. L. K. (2013). Class 3 Semaphorin Mediates Dendrite Growth in Adult Newborn Neurons through Cdk5/FAK Pathway. PLoS ONE, 8(6), 1–15.

42. Nishimura, Y. V, Shikanai, M., Hoshino, M., Ohshima, T., & Nabeshima, Y. (2014). Cdk5 and its substrates, Dcx and p27 kip1, regulate cytoplasmic dilation formation and nuclear elongation in migrating neurons. Development, 141(18), 3540–3550.

43. Nixon, R. A., & Shea, T. B. (1992). Dynamics of neuronal intermediate filaments: A developmental perspective. Cell Motility and the Cytoskeleton, 22(2), 81–91.

44. Ohshima, T., Ward, J. M., Huht, C., Longenecker, G., Veeranna, Pant, H. C., Brady, R., Martin, L.J., Kulkarni, A. B. (1996). Targeted disruption of the cyclin-dependent kinase 5 gene results in abnormal corticogenesis, neuronal pathology and perinatal death. PNAS, 93(October), 11173–11178.

45. Oshima, T., Motoyuki, H., Tabata, H., Mutoh, T., Adachi, T., Hiromi, S., Saruta, K., Iwasato, T., Itohara, S., Hashimoto, M., Nakajima, K., Ogawa, M., Kulkarni, A., Mikoshiba, K. (2007). Cdk5 is required for multi-to-bipolar transition during radial neuronal migration and proper dendrite development of pyramidal neurons in the cerebral cortex. Development, 134, 2273–2282

46. Pallari, H., Lindqvist, J., Torvaldson, E., Ferraris, S. E., & He, T. (2011). Nestin as a regulator of Cdk5 in differentiating myoblasts. Molecular Biology of the Cell, 22, 1539–1549.

47. Park, D., Xiang, A. P., Mao, F. F., Zhang, L., Di, C. G., Liu, X. M., … Lahn, B. T. (2010). Nestin is required for the proper self-renewal of neural stem cells. Stem Cells, 28(12), 2162–2171.

48. Pennypacker, K., Fischer, I., Levitt, P. (1991). Early in Vitro Genesis and Differentiation of Axons and Dendrites by Hippocampal Neurons Analyzed Quantitatively with Neurofilament-H and Microtubule-Associated Protein 2 Antibodies. Experimental Neurology, 111(Jan), 25–35.

49. Petrovic, M., & Schmucker, D. (2015). Axonal wiring in neural development: Target-independent mechanisms help to establish precision and complexity. Bioessays, 37(Sep), 996–1004.

50. Polleux, F., Giger, R., Ginty, D., Kolodkin, A., Chosh, A. (1998). Patterning of Cortical Efferent Projections by Semaphorin-Neuropolin Interactions. Science, 282(5395) 1904–1906

51. Sahlgren, C. M., Mikhailov, A., Vaittinen, S., Pallari, H., Kalimo, H., Pant, H. C., & Eriksson, J. E. (2003). Cdk5 Regulates the Organization of Nestin and Its Association with p35. Molecular and Cellular Biology, 23(14), 5090–5106.

52. Sahlgren, C. M., Pallari, H.-M., He, T., Chou, Y.-H., Goldman, R. D., & Eriksson, J. E. (2006). A nestin scaffold links Cdk5/p35 signaling to oxidant-induced cell death. The EMBO Journal, 25(20), 4808–4819.

53. Sakakibara, A., & Hatanaka, Y. (2015). Neuronal polarization in the developing cerebral cortex. Frontiers in Neuroscience, 9(April), 1–10.

54. Sapir, T., Horesh, D., Caspi, M., Atlas, R., Burgess, H. a, Wolf, S. G., Francis, F., Chelly, J., Elbaum, M., Pietrokovski, S., Reiner, O. (2000). Doublecortin mutations cluster in evolutionarily conserved functional domains. Human Molecular Genetics, 9(5), 703–712.

55. Schaar, B. T., Kinoshita, K., & McConnell, S. K. (2004). Doublecortin Microtubule Affinity Is Regulated by a Balance of Kinase and Phosphatase Activity at the Leading Edge of Migrating Neurons. Neuron, 41(2), 203–213.

56. Shaw, G., Banker, G., Weber, K. (1985). An immunofluorescence study of neurofilament protein expression by developing hippocampal neurons in tissue culture. European Journal of Cell Biology, 39(1), 205–216.

57. Shin, E., Kashiwagi, Y., Kuriu, T., Iwasaki, H., Tanaka, T., Koizumi, H., Gleeson, JG., Okabe, S. (2013). Doublecortin-like kinase enhances dendritic remodelling and negatively regulates synapse maturation. Nature Communications, 4(4), 1440.

58. Shmueli, A., Gdalyahu, A., Sapoznik, S., Sapir, T., Tsukada, M., & Reiner, O. (2006). Site-specific dephosphorylation of doublecortin (DCX) by protein phosphatase 1 (PP1). Molecular and Cellular Neuroscience, 32(1–2), 15–26.

59. Su, P.-H., Chen, C.-C., Chang, Y.-F., Wong, Z.-R., Chang, K.-W., Huang, B.-M., & Yang, H.-Y. (2013). Identification and cytoprotective function of a novel nestin isoform, Nes-S, in dorsal root ganglia neurons. The Journal of Biological Chemistry, 288(12), 8391–8404.

60. Szaro, B. E. N. G., Grant, P., Lee, V. M., & Gainer, H. (1991). Inhibition of Axonal Development After Injection of Neurofilament Antibodies Into a Xenopus Zaeuis Embryo. Journal of Comparative Neurology, 585, 576–585.

61. Tamariz, E., Varela-echavarría, A. (2015). The discovery of the growth cone and its influence on the study of axon guidance. Frontiers in Neuroanatomy, 9(May), 1–9.

62. Tanaka, T., Serneo, F. F., Higgins, C., Gambello, M. J., Wynshaw-Boris, A., & Gleeson, J. G. (2004). Lis1 and doublecortin function with dynein to mediate coupling of the nucleus to the centrosome in neuronal migration. The Journal of Cell Biology, 165(5), 709–721.

63. Tanaka, T., Serneo, F. F., Tseng, H. C., Kulkarni, A. B., Tsai, L. H., & Gleeson, J. G. (2004). Cdk5 Phosphorylation of Doublecortin Ser297 Regulates Its Effect on Neuronal Migration. Neuron, 41(2), 215–227.

64. Tsukada, M., Prokscha, A., & Eichele, G. (2006). Neurabin II mediates doublecortin-dephosphorylation on actin filaments. Biochemical and Biophysical Research Communications, 343(3), 839–847. https://doi.org/10.1016/j.bbrc.2006.03.045

65. Tsukada, M., Prokscha, A., Ungewickell, E., & Eichele, G. (2005). Doublecortin association with actin filaments is regulated by neurabin II. The Journal of Biological Chemistry, 280(3), 11361–11368.

66. Uchida, A., & Brown, A. (2004). Arrival, Reversal, and Departure of Neurofilaments at the Tips of Growing Axons. Molecular Biology of the Cell, 15(September), 4215– 4225.

67. Vitriol, E. A., & Zheng, J. Q. (2012). Growth Cone Travel in Space and Time: The Cellular Ensemble of Cytoskeleton, Adhesion, and Membrane. Neuron, 73(6), 1068–1080.

68. Walker, K. L., Yoo, H. K., Undamatla, J., & Szaro, B. G. (2001). Loss of neurofilaments alters axonal growth dynamics. The Journal of NeuroscienceL 21(24), 9655–9666.

69. Wang, J., Huang, Y., Cai, J., Ke, Q., Xiao, J., Huang, W., Li, H., Qiu, Y., Wang, Y., Shang, B., Wu, H., Zhang, Y., Sui, X., Bardeesi, AS., Xiang, A. (2017). A Nestin – Cyclin-Dependent Kinase 5 – Dynamin-Related Protein 1 Axis Regulates Neural Stem / Progenitor Cell Stemness via a Metabolic Shift. Stem Cells, 36, 589–601.

70. Wang, L., Ho, C., Sun, D., Liem, R. K. H., & Brown, A. (2000). Rapid movement of axonal neurofilaments interrupted by prolonged pauses. Nature Cell Biology, 2(March), 137–141.

71. Wang, X., Sterne, G. R., & Ye, B. (2014). Regulatory mechanisms underlying the differential growth of dendrites and axons. Neuroscience Bulletin, 30(4), 557–568.

72. Wilhelmsson, U., Lebkuechner, I., Leke, R., Marasek, P., Yang, X., Antfolk, D., Chen, M., Mohseni, P., Lasic, E., Bobnar, ST., Stenovec, M., Zorec, R., Nagy, A., Sahlgren, C., Pekna, M., Pekny, M. (2019). Nestin Regulates Neurogenesis in Mice Through Notch Signaling From Astrocytes to Neural Stem Cells. Cerebral Cortex, 1–17.

73. Xie, Z., Samuels, B. A., & Tsai, L. H. (2006). Cyclin-dependent kinase 5 permits efficient cytoskeletal remodeling - A hypothesis on neuronal migration. Cerebral Cortex, 16, 64–68.

74. Yabe, J. T., Chan, W. K. H., Wang, F. S., Pimenta, A., Ortiz, D. D., & Shea, T. B. (2003). Regulation of the Transition From Vimentin to Neurofilaments During Neuronal Differentiation. Cell Motility and the Cytoskeleton, 56(March), 193–205.

75. Yamada, K. M., Spooner, B. S., & Wessells, N. K. (1970). Axon Growth: Roles of Microfilaments and Microtubules. Proceedings of the National Academy of Sciences, 66(4), 1206–1212.

76. Yamamoto, N., Tamada, A., & Murakami, F. (2003). Wiring of the brain by a range of guidance cues. Progress in Neurobiology, 68(Dec), 393–407.

77. Yang, J., Dominguez, B., de Winter, F., Gould, T. W., Eriksson, J. E., & Lee, K.-F. (2011). Nestin negatively regulates postsynaptic differentiation of the neuromuscular synapse. Nature Neuroscience, 14(3), 324–330.

78. Yap, C. C., Digilio, L., Kruczek, K., Roszkowska, M., Fu, X., Liu, J. S., & Winckler, X. B. (2018). A dominant dendrite phenotype caused by the disease-associated G253D mutation in doublecortin (DCX) is not due to its endocytosis defect. Journal of Biological Chemistry, 293(12), 18890–18902.

79. Yap, C. C., Digilio, L., Mcmahon, L., Roszkowska, M., Bott, C. J., Kruczek, K., & Winckler, X. B. (2016). Different Doublecortin (DCX) Patient Alleles Show Distinct Phenotypes in Cultured Neurons. Journal of Biological Chemistry, 291(52), 26613– 26626.

80. Yap, C. C., Vakulenko, M., Kruczek, K., Motamedi, B., Digilio, L., Liu, J. S., & Winckler, B. (2012). Doublecortin (DCX) mediates endocytosis of neurofascin independently of microtubule binding. The Journal of Neuroscience 32(22), 7439–7453.

81. Yuan, A., & Rao, M. V. (2012). Neurofilaments at a glance. J Cell Sci, 125(14), 3257– 3263.

82. Zhang, Y., Wang, J., Huang, W., Cai, J., Ba, J., Wang, Y., Ke, Q., Huang, Y., Liu, X., Qiu, Y., Lu, Q., Sui, X., Shi, Y., Wang, T., Shen, H., Guan, Y., Zhou, Y., Chen, Y., Wang, M., Xiang, A. P. (2018). Nuclear Nestin deficiency drives tumor senescence via lamin A/C-dependent nuclear deformation. Nature Communications, 9(1), 1–14.

